# Proteomic Subtyping of Alzheimer’s Disease CSF links Blood-Brain Barrier Dysfunction to Reduced levels of Tau and Synaptic Biomarkers

**DOI:** 10.1101/2025.03.14.643332

**Authors:** Madison C. Bangs, Joshna Gadhavi, E. Kathleen Carter, Lingyan Ping, Duc M. Duong, Eric B. Dammer, Fang Wu, Anantharaman Shantaraman, Edward J. Fox, Erik C.B. Johnson, James J. Lah, Allan I. Levey, Nicholas T. Seyfried

## Abstract

Alzheimer’s disease (AD) is characterized by significant clinical and molecular heterogeneity, influenced by genetic and demographic factors. Using an unbiased, network-driven approach, we analyzed the cerebrospinal fluid (CSF) proteome from 431 individuals (483 samples), including 111 African American participants, to identify core protein modules associated with AD, race, sex, and age. Our analysis revealed ten co-expression modules linked to distinct biological pathways and cell types, many of which correlated with established AD biomarkers such as β-amyloid, tau, and phosphorylated tau. To further resolve disease heterogeneity, we applied a proteomic subtyping approach, identifying six distinct CSF subtypes spanning the clinical and pathological spectrum. These subtypes were validated across independent cohorts, with many aligning with previously defined AD subtypes, including those linked to neuronal hyperplasticity, immune activation, and blood-brain barrier (BBB) integrity. Notably, the BBB subtype, enriched with African Americans and men, was characterized by low CSF tau, high CSF/serum albumin ratios, and reduced synaptic protein levels. This subtype also exhibited increased levels of proteolytic enzymes, including thrombin and matrix metalloproteases, that cleave tau. Plasma dilution into the neuronal hyperplastic AD subtype CSF led to reduced tau and synaptic protein module levels, indicating that plasma protease activity contributes to tau and synaptic protein depletion independent of underlying brain pathology. These findings highlight the impact of BBB integrity on CSF tau levels, particularly in men and African Americans, and underscore the need for diversity-informed AD biomarker strategies to improve diagnostics and therapeutic targeting across populations.

## Introduction

The neuropathological changes of Alzheimer’s disease (AD) begin two or more decades before signs of cognitive impairment, making early detection crucial for disease prevention and therapeutic engagement^1, 2^. The detection and progression of AD is frequently quantified using the ATN framework, which balances the contributions of the three major axes of neuropathology in AD: production and accumulation of Aβ plaques (A), tau tangles (T), and neurodegeneration (N)^3^. Peripheral biomarkers of amyloid, tau, and phosphorylated tau (pTau) measured in cerebrospinal fluid (CSF) reflect underlying brain pathologies and have predominantly been used to stage disease progression^4^. However, these established biomarkers do not always progress in tandem to manifest a uniform presentation of the disease^5^. Genetic risk factors, sex, race, and environmental exposures can all modulate disease biology through mechanisms that have not yet been fully elucidated. For example, African Americans with AD have lower levels of CSF tau compared to non-Hispanic Whites (NHW)^6, 7^, despite having equivalent levels of tau and amyloid in the brain at autopsy^8, 9^, and a higher lifetime incidence of dementia^10–17^. This disparity may contribute to delayed or under-diagnosis in African Americans^11^. Furthermore, pathological complexity and heterogeneity are the rule, rather than the exception in AD, with approximately 70% of AD dementia cases including additional age-related pathologies at autopsy, such as cerebrovascular disease, Lewy body inclusions, and TAR DNA-binding protein 43 (TDP-43) aggregates^18–21^. However, biomarkers of amyloid and tau do not accurately predict the extent of cognitive impairment, with some cognitively impaired individuals often showing normal amyloid levels, and 30-40% of cognitively unimpaired elderly individuals showing AD pathology^1, 22^. Thus, an integrated approach to resolve disease heterogeneity is required to provide the most effective method of characterizing and potentially treating the differing molecular manifestations of AD.

Molecular subtyping has been proposed to address these issues by providing a more nuanced understanding of the clinical and pathological heterogeneity of AD. Recent efforts have utilized genomic, transcriptomic, proteomic, and metabolomic analyses, which have been pivotal in identifying unique AD subtypes associated with diverse clinical and pathological phenotypes in brain and biofluids^23–27^. Recent evidence from unbiased CSF proteomics approaches indicates that AD has at least five core subtypes that reflect underlying pathological and clinical heterogeneity^26^. These subtypes have been linked to neuronal hyperplasticity, immune system activation, RNA metabolism, choroid plexus function, and blood–brain barrier (BBB) integrity, each following different trajectories, with varying degrees of amyloid, tau, and neurodegeneration biomarkers^26^. Whether these five subtypes can be resolved using orthogonal data-driven approaches remains unclear. Additionally, how these subtypes differ across age, race, and sex has not been fully elucidated, as these studies typically adjust for sex-specific variations from their data prior to analysis and have mainly been from larger predominately European or white cohorts^24–26, 28^. Finally, the mechanisms responsible for driving unique subtype differences, such as those linked to distinct synaptic and blood-brain barrier proteomic profiles, which differ in diagnostic biomarker levels, have not been clarified. Understanding these mechanisms is crucial improving diagnostic accuracy and understanding potential discrepancies in treatment outcomes, such as with anti-amyloid therapies.

Here, we have taken an unbiased network-driven approach to identify core protein modules and characterize the CSF proteome, based not only on disease, but also race, age and sex. With this network-based strategy, we analyzed 431 individuals across 483 samples, including 131 from African American participants, and organized the CSF proteome into 10 co-expression modules linked to diverse molecular functions, pathways, and brain cell types. These modules were shown to be associated with age, sex, and race, as well as core AD biomarkers, including Aβ, tau, and pTau, highlighting interactions between diagnostic or progressive AD biomarkers and an individual’s biological traits. To address this heterogeneity, we further clustered both control and AD CSF samples into proteomic subtypes, identifying six distinct proteomic subtypes spanning the spectrum of clinical and pathological burden. These subtypes were validated across independent proteomics datasets, with many of the AD-enriched subtypes aligning with one or more of the five previously mentioned subtypes. Importantly, we show that both African Americans and males were significantly enriched in the BBB subtype, which was characterized by low overall CSF tau, high CSF serum/albumin ratio and low neuronal/synaptic protein levels. We further show that the BBB subtype is enriched in proteolytic enzymes, including thrombin, plasminogen, and matrix metalloprotease (MMPs), which can directly cleave tau. By diluting plasma into CSF *ex vivo* to mimic BBB dysfunction and breakdown, we demonstrate that with increasing plasma levels there is a corresponding reduction of tau and synaptic proteins in AD CSF, in part through thrombin proteolytic activity. These data suggest that the enzymatic activity of plasma due to BBB breakdown may result in lower tau and synaptic protein levels in CSF, independent of underlying brain pathology, which could reconcile the lower levels of CSF tau observed in African Americans. Thus, the development of replicable and diversity informed CSF subtypes can help enable clearer insight into the driving biological modifiers of AD, allowing for more targeted and effective therapeutic interventions and an enhanced understanding of the factors impacting disease progression.

## Results

### Demographic and AD Biomarker Characterization of a Consensus CSF Proteomic Dataset

We generated a consensus CSF proteomic dataset by harmonizing tandem mass tag-based mass spectrometry (TMT-MS) proteomics across previously published studies^29, 30^. In total, 483 CSF samples (n = 238 AD, n = 245 controls) were analyzed, sourced from Emory’s Goizueta Alzheimer’s Disease Research Center (GADRC) (**Fig. 1A**). These included samples from two sets: Set 1 (n = 281) and Set 2 (n = 202), with 52 technical replicates between them. Broken out demographically, 131 samples were from self-identified African Americans (AA), and 352 from Non-Hispanic Whites (NHW), 178 samples from men, and 305 from women (**Supplemental Table 1**). This combined Emory cohort was searched against the human protein database, identifying and quantifying 2,067 proteins in greater than 50% of samples. Set and batch effects were removed from the data using regression^31^, while preserving the effects of race, sex, and age on protein variation to accurately capture potential contributors to disease heterogeneity (**Supplemental Figure 1A**). Diagnostic biomarkers Aβ_42_, total tau (tTau), and phosphoTau_181_ (pTau_181_) were measured by immunoassay, and cognitive function was assessed using the Montreal Cognitive Assessment (MoCA)^32^. AD diagnosis was determined through consensus assessment at the Emory GADRC, with all AD cases having a tTau/Aβ_42_ immunoassay ratio ≥ 0.226. As immunoassays were run on separate platforms between sets (Set 1: Luminex, Set 2: Roche Elecsys), z-scores were utilized for inter-set comparisons (**Supplemental Table 2**). The strong correlation between immunoassay tTau and independently assessed proteomic measurements of Tau (MAPT) (cor = 0.79, p = 2.9e-104) validates the robustness of the proteomic data and highlights the reliability of using both approaches to study Tau-related processes in the CSF (**Fig. 1B**). Previous studies have demonstrated that across AD cases, levels of CSF Tau are lower in African Americans compared to Non-Hispanic Whites^6, 7, 30^. Consistent with these studies, our analysis showed that levels of both total and pTau were significantly lower in AAs across AD participants (Tukey HSD corrected 3-way ANOVA, tTau: 3.35e-5, pTau: 0.0012) (**Fig. 1C-D**)(**Supplemental Tables 3-4**), However, in our dataset, this trend was driven predominately by African American men. While levels of tTau were significantly lower in all males with AD (Tukey HSD corrected 3-way ANOVA, AD males vs AD females: 1.40e-3), African American men with AD had significantly lower levels of tTau compared to both male and female NHW AD groups (AD-AA males vs AD-NHW males: 0.0015, and females: 1.86e-7), and were trending when compared to their female AA counterparts (p value= 0.052) (**Fig. 1C**). To a lesser extent, these trends were also observed with pTau (AD-AA males vs AD-NHW males: 0.0035, and females: 2.74e-4) (**Fig. 1D**). Within the AD group, Aβ does not differ statistically between race or sex (**Fig. 1E**). However, across all members of the cohort, significant differences were seen in both race and sex, with higher levels of CSF Aβ observed in females and NHWs (Tukey HSD corrected 3-way ANOVA, Sex: 0.0103, Race: 0.0024). The significant variations observed in immunoassay Tau levels when sex and race are broken out in AD cases demonstrates that demographic factors are likely important drivers of disease heterogeneity. The presence of similar trends in other key AD proteins (**Supplemental Fig. 1B**) indicates that the significant and intersecting effects of sex and race seen on tau levels may also influence other pathological pathways, contributing to disease heterogeneity at a molecular level.

**Figure 1:**
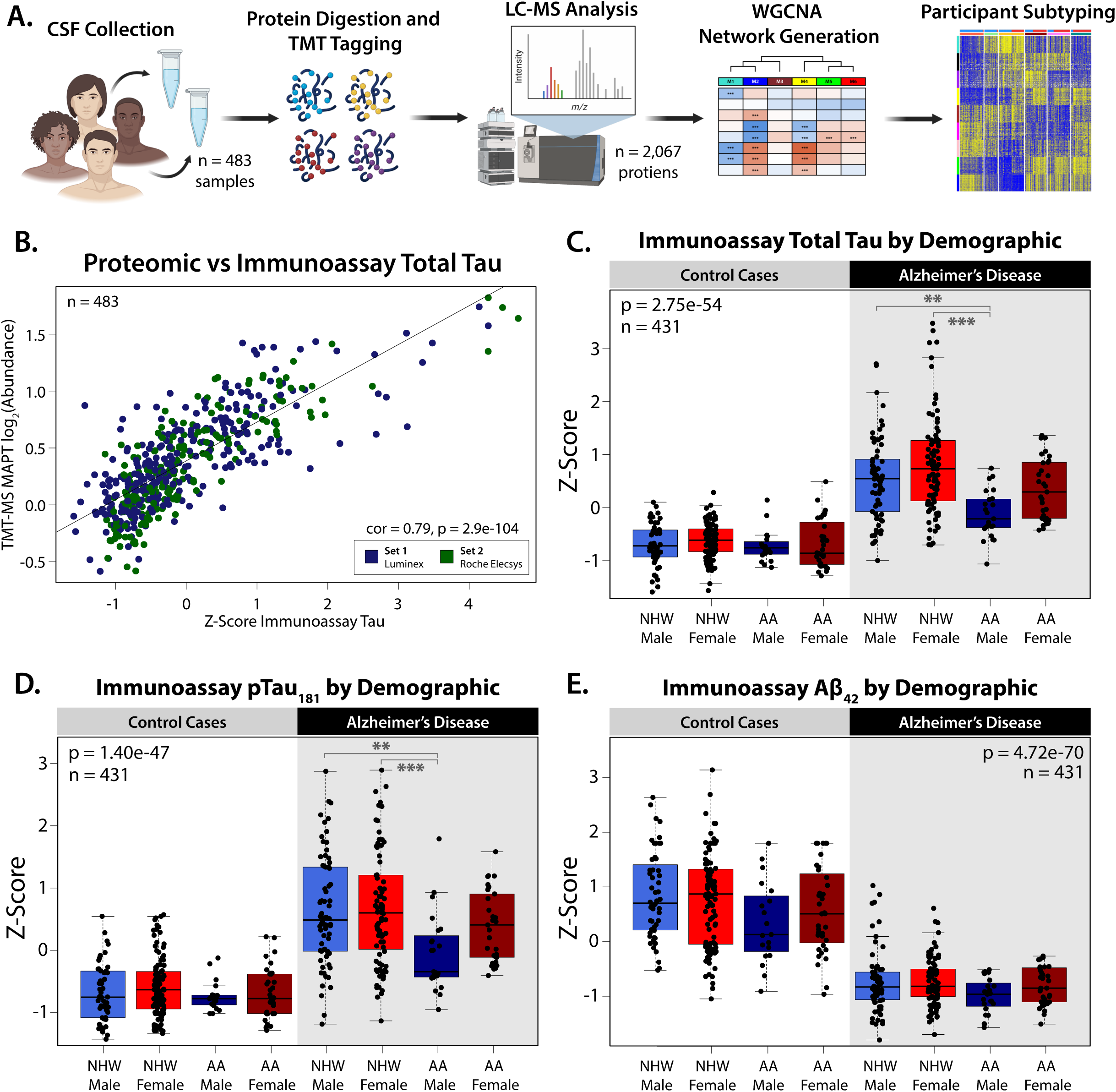
Demographic and Biomarker Characterization of a Consensus CSF Proteomic Dataset Across Race and Sex. **(A)** Study schematic outlining the sample collection, processing, and analysis for subtyping the cerebrospinal fluid (CSF) proteome. **(B)** Immunoassay of total Tau z-scored across Set 1 (Luminex, n = 281) and Set 2 (Roche Elecsys n = 202) was significantly correlated with log_2_ abundance of TMT-MS quantified MAPT (cor = 0.79, p = 2.9e-104). **(C)** Levels of z-score immunoassay total Tau (1-Way ANOVA: 2.75e-54), **(D)** phospho-Tau_181_ (1-Way ANOVA: 1.40e-47), and **(E)** Aβ_42_ (1-Way ANOVA: 4.72e-70). Samples from participants were grouped by diagnosis (AD: 218, Control: 213) and stratified by sex (Male: 164, Female: 267), and self-identified race (Non-Hispanic White (NHW): 320, African American (AA): 111). In same-case sample pairs (n=52), only cases from Set 1 were plotted and included in statistical analysis. Additionally, only points within 3 standard deviations of each group are plotted. Significance for comparison between groups was assessed by three-way ANOVA, across Race, Sex, and Diagnosis, with Tukey post-hoc adjustment (*p≤0.05; **p≤0.01; ***p≤0.001, ****p≤0.0001).

### Network-Based Proteomic Analysis Reveals Cell-Type and Demographic Influences on AD Biomarkers

To investigate the impact of biological traits such as age, sex, and race on the broader AD CSF proteome, we employed an unbiased network-based approach using Weighted Co-expression Network Analysis (WGCNA). This method clusters proteins based on shared expression patterns across individual samples, grouping them into biologically meaningful modules that reflect demographic traits (age, sex and race), disease status, and cell-type origins^33–35^. The 2,067 proteins measured in CSF were organized into 10 distinct modules (M1–M10) and each module was annotated based on Gene Ontology (GO) enrichment and cell-type markers to infer biological processes and brain cell-type origins (**Fig. 2A**) (**Supplemental Tables 6-9**). Additionally, eigenproteins, which represent the first principal component of each module, were correlated with demographic variables (age, sex, and race) and clinicopathological traits (immunoassay levels of Aβ, Tau, pTau, MoCA scores, and APOE genotype) to uncover key proteomic patterns.

**Figure 2:**
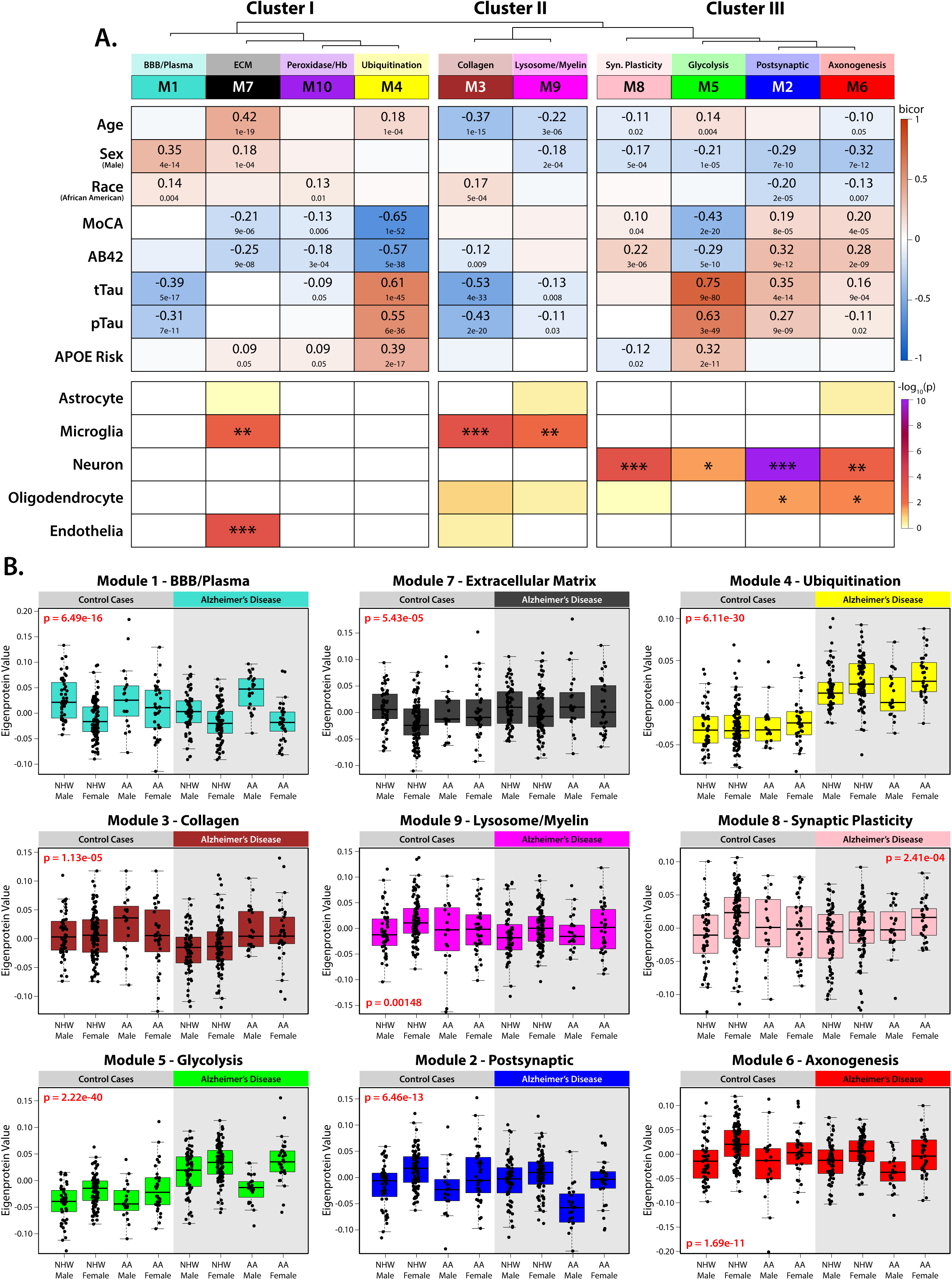
Unbiased network analysis groups the CSF proteome into biologically related modules, which correlate with demographic and clinicopathological traits. **(A)** The Weighted Gene Co-expression Network Algorithm (WGCNA) was used to construct a protein network based on shared cellular expression and biology. This network comprised 10 protein modules and is represented with a dendrogram indicating module relatedness. Gene Ontology (GO) analysis was performed to identify the driving biology represented by each module. Module-trait associations were determined by calculating correlations (bicor) between participant eigenproteins for each module to traits across demographics (age, sex, race), cognitive scores (MoCA) and pathophysiological markers such as immunoassay amyloid beta (Aβ_42_), total Tau (tTau), phospho-Tau_181_ (pTau), and APOE risk score. These module-trait correlations indicate protein groups with strong positive (red) or negative (blue) relationships to participant and disease traits. For same-case sample pairs, only samples from Set 1 were used in calculations. The cell type enrichment of each module was determined by one-tailed Fisher’s exact test (*p≤0.05; **p≤0.01; ***p≤0.001). **(B)** Module eigenprotein values broken out by demographic groups for significant modules, as assessed by one-way ANOVA. Points outside of 3 standard deviations for each demographic group were not plotted.

Network analysis revealed that CSF protein modules were organized into three major clusters based on relatedness (**Fig. 2A**). Cluster I contained four modules, three of which (M1, M10 and M4) had no significant enrichment for a specific brain cell type, however, all were associated with CSF AD biomarkers (Aβ, total tau, and pTau_181_) and demographic traits. Within this cluster, M4, the Ubiquitination module, showed strong correlations with CSF Aβ, total tau, and pTau_181_, as well as APOE genetic risk (quantified by number of the APOE2, APOE3 and APOE4 alleles, scaled by risk) and cognitive impairment (assessed via MoCA score). This module included key neurodegeneration markers such as 14-3-3 family proteins (YWHAE, YWHAZ), neurofilament proteins (NEFL, NEFM), and inflammatory markers (ITGB2, MIF, LDHA). M1, the Blood-Brain Barrier (BBB)/Plasma module, contained plasma-derived proteins, including albumin, immunoglobulins, and thrombin (F2), and was negatively correlated with tau biomarkers. This module was significantly positively correlated with race (increased in African Americans) and sex (increased in males). M10, the Peroxidase/Hemoglobin module, was comprised of antioxidant-associated proteins, including peroxisomal proteins (CA1, CA2, CAT) and peroxiredoxins (PRDX6, PRDX2). Although it contained hemoglobin-related proteins, it did not share the demographic associations observed in M1. Furthermore, eigenproteins of each module were not strongly correlated across samples (bicor = 0.23, p = 1e-6) (**Supplemental Figure 2**). The final Cluster I module, M7, the Extracellular Matrix module, was the only module associated with brain cell type signatures and was enriched for microglial and endothelial markers. This module showed higher abundance in males and was positively associated with age, and to a lesser degree cognitive decline (increasing with age and with worsening cognitive deficits by MoCA).

Cluster II contained microglia-associated modules, including M3 (Collagen) and M9 (Lysosome/Myelination). Unlike the microglial module M7 in Cluster I, both M3 and M9 were negatively correlated with age, with lower module levels in older individuals. Both modules also displayed negative correlations with tTau and pTau, potentially representing associations with different microglial activation states or inflammatory responses compared to M7. Levels of M3 (Collagen) were increased in African Americans, while levels of M9 (Lysosome/Myelin) were higher in women.

Cluster III consisted of neuron-enriched modules, including M8 (Synaptic Plasticity), M5 (Glycolysis), M2 (Postsynaptic), and M6 (Axonogenesis). Each of these modules had a significant positive correlation with sex, all with higher abundances in females. To a lesser degree, M2 and M6 were also enriched for oligodendrocyte markers and were significantly lower in African Americans. M5, the Glycolysis module, was primarily composed of neuronally associated hallmarks of AD, such as tau (MAPT), NRGN, GAP43, CALM1, CAMK2A, and SYN1, but also included other AD proteins such as the plaque associated proteins SMOC1 and SPON1^36, 37^. Not surprisingly, M5 was strongly positively correlated with all AD biomarkers and APOE4 genotype, but unlike M4, it was also negatively correlated with sex, with higher levels in females. In contrast to the pathological signature of M5, the other modules in Cluster III, M2, M6, and M8, were associated with cognitive preservation. These modules were positively correlated with higher MoCA scores and lower levels of Aβ in the CSF. M2 contained several key proteins previously linked to cognitive resilience in AD^28^, such as VGF, BDNF, NPY, and NRN1^38, 39^, markers that have also previously been shown to be lower in the CSF of African Americans and decrease with AD progression^30^. M6 (Axonogenesis) contained proteins associated with neuroprotection, such as NPTX2, while M8 (Synaptic plasticity) was the only module in the network with a negative correlation to APOE4 risk.

All 10 protein modules were significantly associated with at least one demographic factor, emphasizing the strong influence of sex, race, and age on the CSF proteome (**Fig 2A-B**). Notably, sex emerged as a major driver of module variance, with strong inverse correlations observed between the two largest modules: M1 (BBB/Plasma), which was significantly elevated in males, and M2 (Postsynaptic), which was significantly higher in females. These modules also showed significant race-related associations, with M1 being increased in African Americans and M2 being increased in Whites, consistent with previous findings^30^. As a result, the combined effects of sex and race were most pronounced in African American males, who showed the highest levels of M1 and the lowest levels of M2, while the opposite pattern was observed in White females. These differences became even more pronounced in AD cases (**Fig. 2B**). Regression analyses adjusting for disease status, sex, and race individually revealed that M1 (BBB/Plasma) and M2 (Postsynaptic) exhibited opposing trends across race and sex, independent of clinical diagnosis. Specifically, M1 was elevated in males and African Americans, while M2 showed the opposite pattern, further reinforcing our network-based findings. (**Supplemental Figure 3**). These results align with previous studies showing that CSF albumin levels are consistently higher in males than females across all ages, though racial differences in this context had not been previously explored^40^. Collectively, this indicates that race and sex play a significant role in shaping the CSF proteome and influencing disease-related pathways. These results highlight the importance of demographic-stratified analyses in AD CSF to better understand disease heterogeneity and enhance biomarker discovery.

### Unbiased Proteomic Subtyping Reveals Distinct CSF Profiles in AD

AD is clinically and pathologically heterogeneous, underscoring the need for classification or subtyping methods that better capture its underlying biology and relatedness across individuals. To generate unbiased proteomic subtypes in the CSF, we applied the MONET M1 clustering method, which groups samples based on protein-level similarities while optimizing modularity between clusters, consistent with the approach previously used for subtyping the brain proteome^24^. Since the protein modules in our CSF network capture a significant proportion of the variance in protein abundance, reflecting distinct biological processes and cell types, we selected the top 30 proteins from each of the 10 modules to classify samples into subtypes. Each sample was then assigned to one of six distinct proteomic subtypes (**Fig. 3A**). Of the 52 same-case sample pairs across Set 1 and Set 2, 46 (88%) were assigned to the same subtype, demonstrating high reproducibility despite samples being processed separately and analyzed on different mass spectrometers. Non-consistent samples were generally not “hub” samples within their MONET M1 assigned subtype and showed low correlation with their respective clusters. Sample pairs that failed to replicate subtype assignments were, therefore, excluded from further analysis. Ultimately, 425 unique individuals were retained in the final subtyping framework.

**Figure 3:**
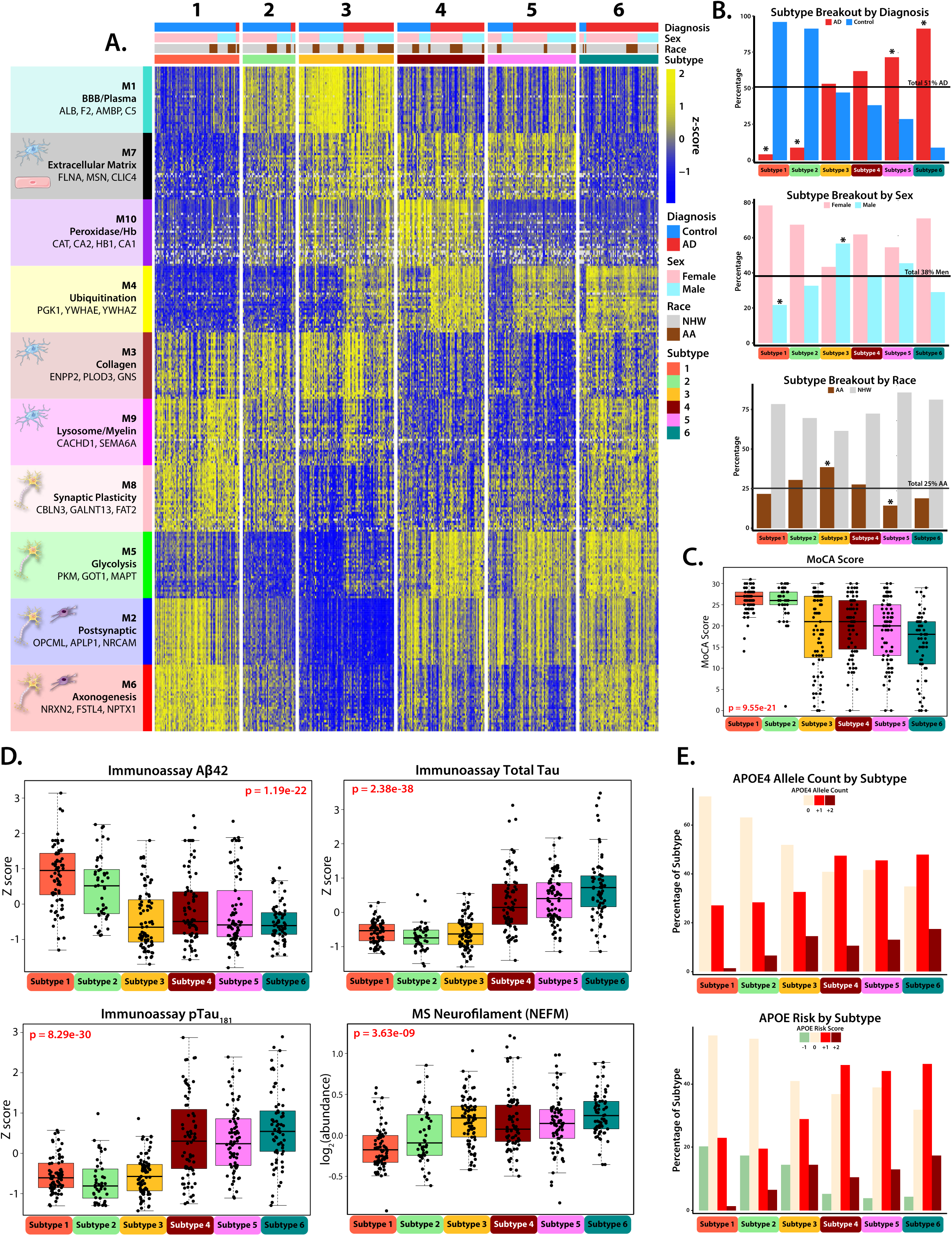
Unbiased CSF subtyping reveals biological heterogeneity in AD, identifying six distinct proteomic subtypes. **(A)** A Heatmap of z-scored abundance of the top 30 proteins from each network module, plotted across participants (n = 425) clustered into the 6 subtypes by the Monet M1 algorithm, with increased protein abundance in yellow, and decreased protein abundance in blue. Individuals who were successfully clustered by the algorithm fell into one of six subtypes (S1: n = 74, S2: n = 46, S3: n = 83, S4: n = 76, S5: n = 77, S6: n = 69). **(B)** The percentage of participants in each subtype compared to the overall demographic percentage within the cohort, broken out by diagnosis (51% AD), sex (38% Male), and self-identified race (25% AA). Subtypes significantly differing from these values as assessed by chi squared test are indicated (*p≤0.05). **(C)** Montreal Cognitive Assessment (MoCA) Score for the participants in each subtype. Significance was assessed by 1-way ANOVA. **(D)** Clinicopathological traits (immunoassay Aβ, tTau, pTau1_181_, and MS Neurofilament (NEFM)) broken out by subtype. Significance was assessed by 1-way ANOVA, points outside of 3 standard deviations for each subtype were not plotted. **(E)** The percentage of each subtype with 0, +1, or +2 copies of the Apolipoprotein E (ApoE) 4 allele (**top**), and ApoE risk score (**bottom**), calculated by adding (+1) for each ApoE4 Allele, and (-1) for each ApoE2 allele, as determined by genetic profiling.

Of the six generated subtypes, Subtypes 1 and 2 were classified as “control-like” because they primarily consisted of clinically normal individuals (S1: n = 74, 96% control; S2: n = 46, 91% control) (**Fig. 3B**). These subtypes exhibited cognitive and biomarker profiles typical of control cases, including low levels of CSF total tau and pTau, higher CSF Aβ, and higher MoCA scores (**Fig. 3C-D**). In contrast, Subtypes 4, 5, and 6 were classified as “AD-like” due to their higher proportion of clinically diagnosed AD participants (S4: n = 76, 62% AD; S5: n = 77, 71% AD; S6: n = 69, 91% AD). All three AD enriched subtypes exhibited CSF biomarker profiles typical of AD, characterized by high levels of CSF total tau and pTau, lower Aβ, and lower MoCA scores (**Supplemental Tables 13-14**).

As expected, when compared to control-like subtypes (S1 and S2), the AD-like subtypes (S4, S5, and S6) had higher levels of M5 (Glycolysis/neuronal) and M4 (Ubiquitination) which have the strongest correlations to tau and amyloid biomarkers in CSF (**Fig. 4**). AD-like Subtypes 4 and 5 exhibited similar proteomic module profiles but were differentiated by the M10 (Peroxidase/Hb) module, which was significantly higher in Subtype 4 than in Subtypes 5 and 6 (**Supplemental Tables 15-16**). Subtype 6 differed from the other two AD-like subtypes by displaying elevated levels of neuronal modules M6 (Axonogenesis) and M8 (Synaptic Plasticity) in Cluster III. It also showed distinct patterns in microglia-associated modules, with higher levels of M9 (Lysosome/Myelination) and lower levels of M7 (Extracellular Matrix) compared to AD-like Subtypes 4 and 5. All AD-like subtypes (S4, S5, and S6) had a higher prevalence of the APOE ε4 genotype compared to the control-like subtypes (S1 and S2), however, no significant differences were found in the prevalence of APOE ε4 among the AD-like subtypes (**Fig. 3E**).

**Figure 4:**
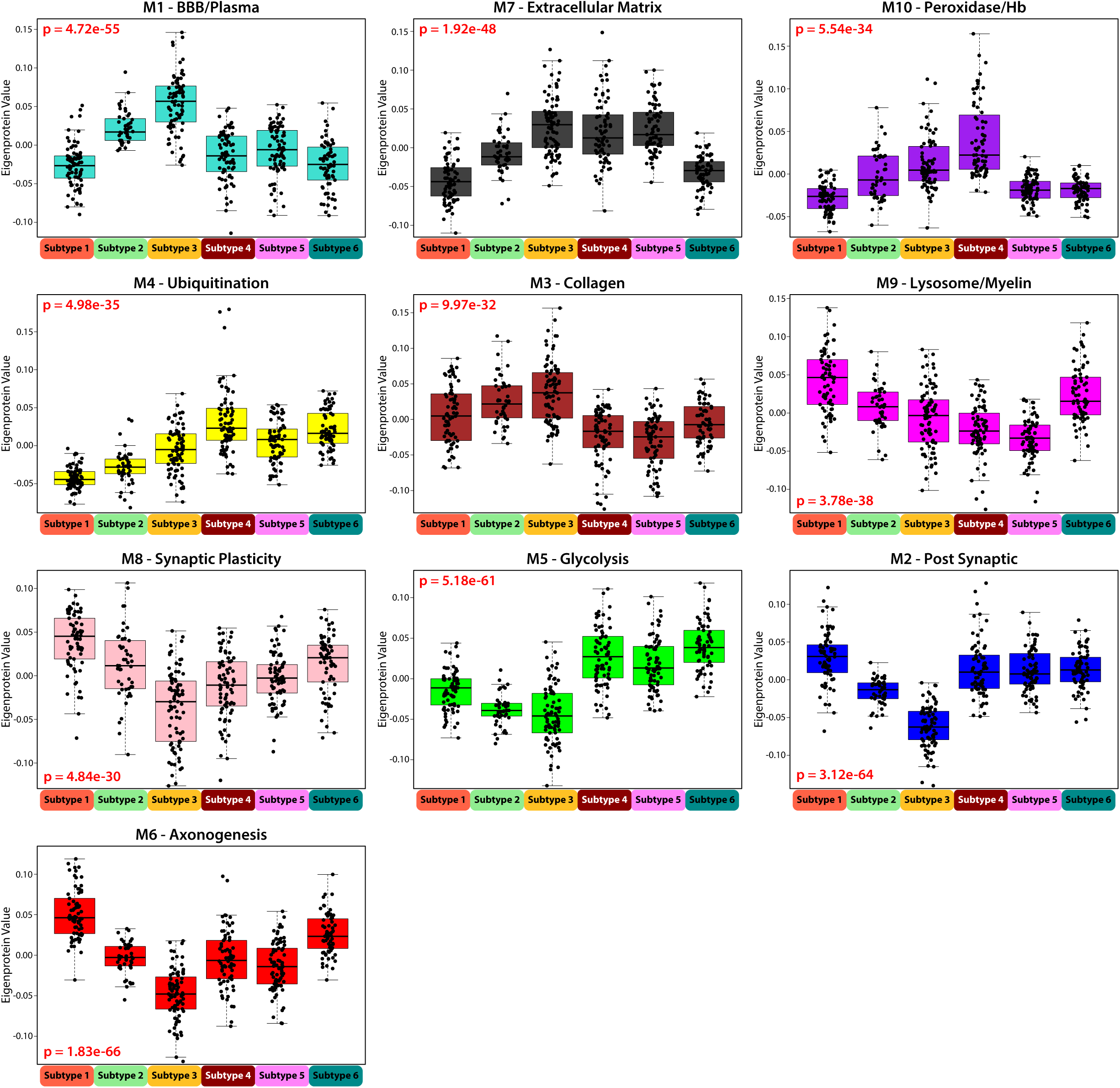
Network Module Eigenprotein Patterns Differentiate Emory Subtypes. Eigenprotein values broken out by network module across the 6 subtypes organized in module relatedness order. Plots 1-5 demonstrate increasing protein abundance across “control-like” Subtypes 1-3, and plots 6-10 demonstrate decreasing protein abundance across Subtypes 1-3. Significance was assessed by 1-way ANOVA, and points outside of 3 standard deviations for each subtype were not plotted.

Subtype 3 was unique in that it was nearly equally comprised of control and AD cases (S3: n = 83, 53% AD). It was distinguished from other subtypes by its distinct proteomic signature, characterized by the highest levels of M1 (BBB/Plasma) across all subtypes and the lowest levels for the Cluster III neuronal modules (M8: Synaptic Plasticity, M5: Glycolysis, M2: Postsynaptic, and M6: Axonogenesis) (**Fig. 4**). This subtype also displayed a unique AD biomarker profile. While individuals in Subtype 3 had lower immunoassay Aβ levels comparable to those seen in AD-like subtypes, their total tau and pTau levels were more like those of the control-like subtypes (S1 and S2). Additionally, members of Subtype 3 exhibited high levels of CSF neurofilament, a marker of neurodegeneration^41^, along with lower average MoCA scores than control-like subtypes. The lower tau levels observed in participants of this subtype are inconsistent with their “AD-like” levels of Aβ and markers of neurodegeneration (NEFL/M and cognitive score). Thus, this subtype represents an atypical (A+/T-) presentation of AD, as elevated tau is usually considered a precursor to neurodegeneration in the ATN framework^42^. Subtype 3 was also significantly enriched for African Americans and males, who exhibited lower immunoassay CSF tau levels in AD cases, which is consistent with the sex and race-based associations observed for M1 (BBB/plasma) and lower M2 (synaptic) for these individuals (**Figs. 1** and **2**). In line with this, the synaptic-enriched M5 (Glycolysis) module, which is strongly correlated with disease pathology and includes tau (MAPT) as a hub protein, was also low in Subtype 3 compared to other AD-like subtypes, suggesting that the lower immunoassay tau levels may be associated with the overall reduction of neuronal proteins in this module. Notably, despite half of Subtype 3 consisting of clinically defined control participants, its distinct proteomic profile was sufficient to cluster these individuals into a unique subtype, whereas the clustering of other subtypes was largely reflective of diagnosis. This distinction was particularly evident in the M4 (Ubiquitination) module, where a clear separation between AD and control cases was observed within Subtype 3 (**Fig. 3A**). In summary, our unbiased proteomic subtyping approach identified six distinct CSF subtypes, including two control-like and three AD-like subtypes that aligned with expected biomarker and network-module profiles. Interestingly, Subtype 3 emerged as an outlier with an atypical A+/T-CSF biomarker profile, characterized by low tau levels despite elevated neurofilament, enrichment in African Americans and men, and distinct proteomic signatures that challenge conventional AD diagnostic classification.

### Validation of CSF Proteomic Subtypes in an Independent Cohort Supports Differences in Established AD Biomarkers and Links to Blood-brain-barrier Dysfunction

To confirm our subtyping results in the Emory cohort, we analyzed a CSF proteomic dataset from a previously published TMT-MS study of 120 community-dwelling participants in Switzerland, including 42 AD patients and 78 cognitively normal individuals^43^ (**Fig. 5A**). In this Swiss CSF dataset, 697 proteins were quantified with greater than 50% completeness across all samples. To assign these Swiss CSF samples to the Emory subtypes, we first generated a supervised Uniform Manifold Approximation and Projection (UMAP) embedding for the 425 Emory samples, using the overlapping proteins (n = 167) as features and the six MONET M1 subtypes as target labels (**Fig. 5B**), which clearly distinguished the six subgroups. When the Swiss samples were projected onto this UMAP space, 115 out of 120 cases mapped to one of the predefined Emory subtypes.

**Figure 5:**
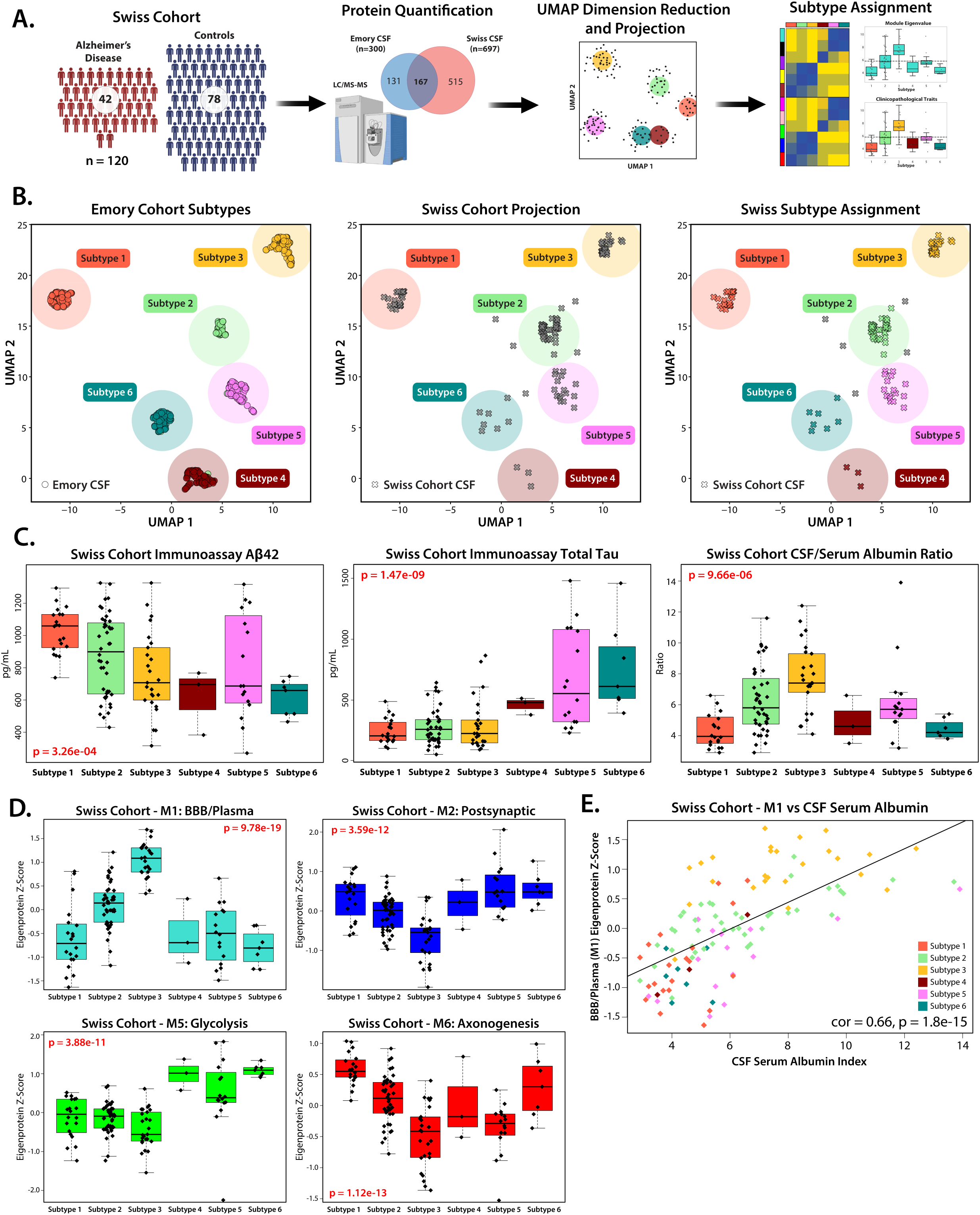
Replication of Emory CSF Subtypes in the Swiss Cohort Using UMAP Projection and AD Biomarker Validation. **(A)** Schematic of project workflow for the assignment of cases in a replication cohort (Swiss Cohort, n=120), via Uniform Manifold Approximation and Projection (UMAP). **(B)** A supervised UMAP embedding was generated (**left**) using overlapping proteins from the Swiss and Emory cohort module hubs (log_2_ abundance, n=167 proteins). Mapping the Emory Cases onto the UMAP dimensions produced with the following settings successfully replicated the original six subtypes in 98.8% of cases: n_neighbors = 10, n_components = 2, metric = Euclidean, and min_dist = 0.1. Cases from the Swiss Cohort were then projected (**center**) and 96% of cases were assigned to one of the six subtypes (**right**). **(C)** Trends in the immunoassay diagnostic biomarkers Aβ (**left**) and Tau (**center**) from the Swiss replication cohort mirror those of the original Emory cohort. CSF/Serum albumin ratio (**right**), a clinical marker of blood brain barrier breakdown, is highest in the replication cohort cases assigned to Subtype 3. Significance was assessed by 1-way ANOVA, points outside of 3 standard deviations for each subtype were not plotted. **(D)** Z-score eigenprotein values for the overlapping proteins in key modules M1, M2, M5 and M6 replicate the trends seen in the Emory cohort. Significance was assessed by 1-way ANOVA, points outside of 3 standard deviations for each subtype were not plotted. **(E)** Levels of M1 proteins were highly correlated with CSF Serum Albumin index (Pearson cor = 0.66, p = 1.8e-15) across all subtypes in the Swiss replication cohort.

While proteins from each module were represented in the overlap, some modules, such as M7 (Extracellular Matrix) and M4 (Ubiquitination), had lower representation (<40% overlap), while M10 (Peroxidase/Hb) had very low representation (<25% overlap), potentially introducing ambiguity in case assignment. Despite these differences in proteome coverage, the clinicopathological traits and module abundances of the Swiss cohort closely mirrored those observed in the Emory subtypes. Specifically, Swiss individuals that mapped to Subtype 3 exhibited low Aβ levels, consistent with AD-like Subtypes 4, 5, and 6, while their tau levels aligned more closely with the control-like Subtypes 1 and 2 (**Fig. 5C**). Additionally, samples in the Swiss cohort that mapped to Subtype 3 displayed the highest levels of M1 (BBB/Plasma) and the lowest levels of Cluster III neuronal protein modules, M2, M6 and M8 (**Fig. 5D and Supplemental Figure 4**). Biological sex data was not available for the Swiss cohort, preventing confirmation of the male enrichment observed in Subtype 3 in the Emory cohort.

Importantly, the Swiss cohort also included CSF/serum albumin ratio measurements for each participant as clinical trait, a biomarker of BBB dysfunction^44, 45^. Individuals classified as Subtype 3 exhibited the highest CSF/serum albumin ratios (**Fig. 5C**), which were strongly correlated with M1 abundance (cor = 0.66, p = 1.8e-15) (**Fig. 5E**), suggesting a potential link between elevated M1 levels in CSF and clinical markers of BBB breakdown^25, 26^. The successful replication of the six CSF proteomic subtypes in this independent European cohort, despite differences in proteome coverage, validates their biological relevance and further supports the link between Subtype 3, blood-brain barrier dysfunction, and distinct tau and Aβ profiles.

### Cross-Cohort Validation of CSF Proteomic Subtypes Reveals Strong Molecular Concordance with Independent AD Subtypes

Recently, Tijms et al. used TMT-MS CSF proteomics to characterize AD heterogeneity, identifying five distinct AD subtypes defined by proteins enriched in neuronal hyperplasticity, innate immune activation, blood-brain barrier dysfunction, RNA dysregulation, and choroid plexus dysfunction, each associated with specific genetic risk factors, cortical atrophy patterns, and survival outcomes^26^. Their study classified only AD cases (n = 419) from the Alzheimer Center Amsterdam (ACA) based on differentially abundant CSF proteins (n = 1,087) compared to cognitively normal individuals (n = 187).

To compare our results with these subtypes, we used Emory Subtype 1, which consists of >95% cognitively normal control individuals, as a reference to determine correlations between the remaining five Emory subtypes and those five AD subtypes identified in the ACA. This was done using a mean-centered z-score per protein, per subtype (**Supplemental Fig. 5**). Each Emory subtype showed a significant positive correlation with at least one AD subtype in the ACA cohort. Consistent with prior hypotheses, Emory Subtype 3 was highly correlated with both the choroid plexus dysfunction (bicor = 0.67, p = 5.76e-39) and blood-brain barrier dysfunction (bicor = 0.79, p = 4.13e-63) AD subtypes in the ACA. Despite being composed primarily of control cases, Emory Subtype 2 also correlated with choroid plexus dysfunction subtype, though to a lesser extent (bicor = 0.23, p = 7.27e-5). Notably, these two subtypes in the ACA exhibited the lowest levels of immunoassay tau, mirroring trends observed in the Emory and Swiss cohorts (**Figs. 3** and **5**). Emory Subtype 4 was most strongly correlated with the innate immune activation subtype (bicor = 0.73, p = 5.13e-49), while Subtype 5 aligned closely with the RNA dysregulation subtype (bicor = 0.79, p = 4.66e-62), though some overlap was observed between the two. The highest correlation was between Emory Subtype 6 and the neuronal hyperplasticity subtype (bicor = 0.88, p = 9.11e-94). Additionally, neuronal hyperplasticity and blood-brain barrier dysfunction subtypes were negatively correlated in the ACA cohort, mirroring the inverse relationship observed between Subtype 6 and Subtype 3 in the Emory Cohort. The strong correlations observed between these subtypes, despite differences in protein selection, patient populations, and clustering methods, suggest a high degree of molecular preservation and reinforce the robustness of these AD subtypes across independent cohorts.

### Blood-Brain Barrier Dysfunction and Protease Activity in Subtype 3 Contribute to Tau Cleavage

Given its central role in defining Subtype 3 and its associations with BBB breakdown, the M1 module (BBB/Plasma) served as a key focal point for investigating biological differences between subtypes. This module consists primarily of plasma-derived proteins, such as albumin, fibrinogen, and immunoglobulins, which are not significantly produced in the brain under normal conditions^46^. Subtype 3, with high M1 levels, was strongly linked to clinical BBB dysfunction metrics by high CSF/serum albumin ratio, suggesting that elevated M1 levels are indicative of BBB impairment regardless of AD status. Across all subtypes, M1 also showed a strong negative correlation with neuronal modules in Cluster III (**Fig. 6A**) and with individual neuronal proteins, including tau (**Fig. 6C**).

**Figure 6:**
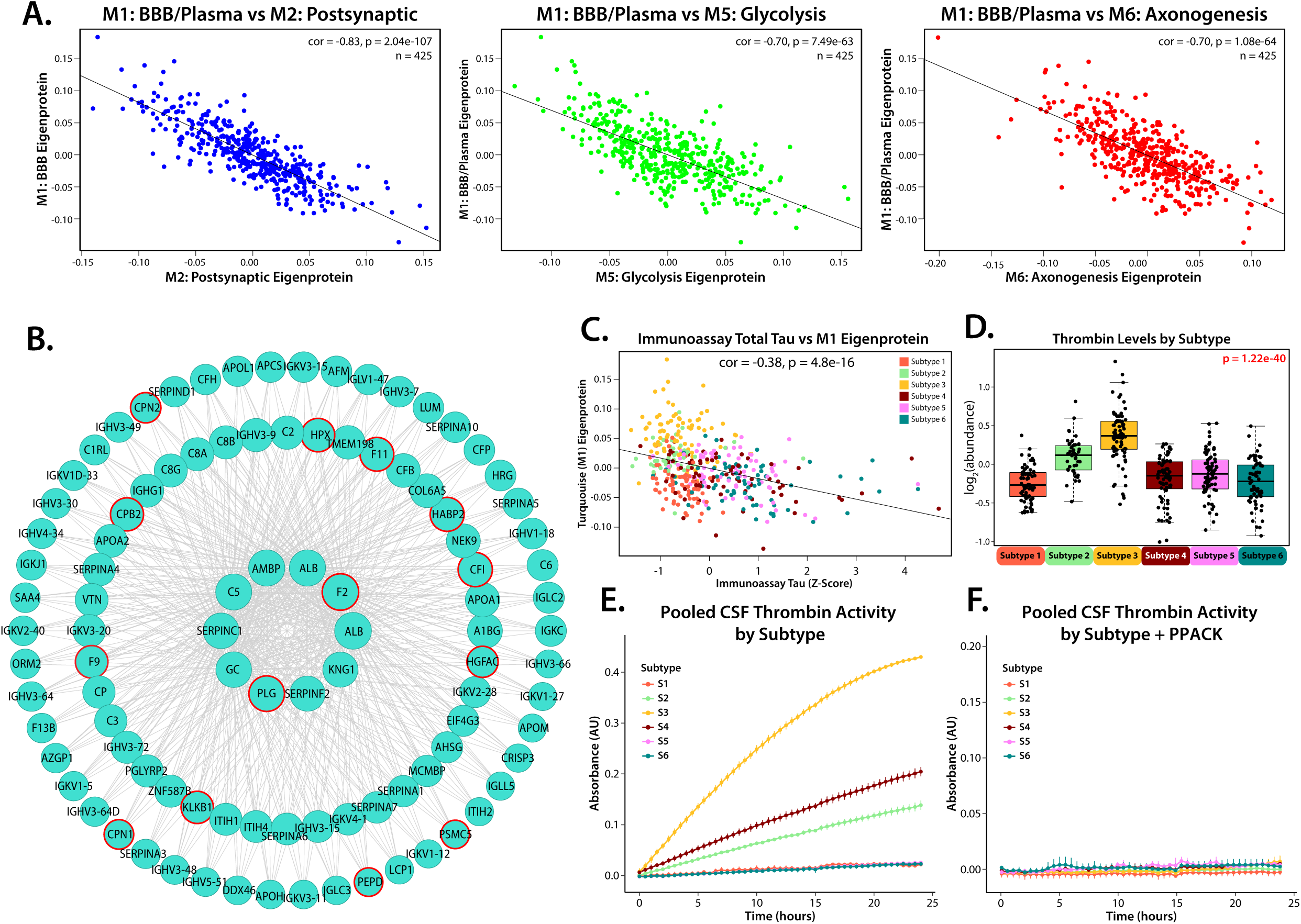
Higher levels of M1, which contains active plasma-derived proteolytic enzymes like thrombin, are strongly correlated with lower levels of CSF Tau and reduced abundance of proteins from neuronal modules. **(A)** Levels of neuronal modules are strongly negatively correlated with levels of M1 (M2: cor = -0.83, p = 2.04 e-107; M5: cor = 0.70, p = 7.49 e-63; M6: cor = -0.70, p = 1.08 e-64) across all subtyped cases. **(B)** Representation of M1 hub proteins including thrombin (F2), produced using igraph, with plasma derived proteolytic enzymes identified by red outline. **(C)** CSF immunoassay total Tau levels are inversely correlated (cor = -0.37, p = 4.1e-17) to levels of M1. **(D)** Levels of thrombin, a plasma derived protein capable of cleaving tau, are highest in Subtype 3. Significance assessed by 1-way ANOVA, points outside of 3 standard deviations for each subtype were not plotted. **(E)** Thrombin activity within pooled samples from each subtype was assessed by thrombin-specific cleavage-based fluorescence assay over 24 hours. **(F)** Thrombin-specific cleavage activity of the pooled CSF samples was inhibited by the addition of the thrombin specific inhibitor PPACK.

Further examination of M1 revealed a significant enrichment of plasma proteases, (such as PLG, F9, MASP2, MMP-2, ADAM10) which account for over 10% of its protein members (**Fig. 6B**). A key hub protein in this module is thrombin (F2), a protease capable of cleaving tau in brain^47^, which was most abundant in the CSF of participants in Subtype 3 (**Fig. 6D**). Thrombin activity, assessed using a thrombin-specific substrate across pooled CSF samples from all six subtypes, demonstrated sustained activity up to 24 hours, with the highest activity observed in Subtype 3 (**Fig. 6E**). The addition of D-Phenylalanyl-L-prolyl-L-arginine chloromethyl ketone (PPACK), a selective and irreversible thrombin inhibitor, successfully mitigated this activity (**Fig. 5F**). These findings indicate that thrombin in Subtype 3 remains in its active form, supporting a potential mechanistic link between high M1 protease levels and reduced tau and synaptic proteins levels in modules mapping to Cluster III.

To explore this hypothesis, we show that recombinant thrombin can directly cleave recombinant biotinylated tau into distinct fragments *in vitro* (**Fig. 7A**). We next investigated whether similar proteolytic activity occurred in plasma and CSF samples from study participants. When recombinant biotinylated 2N4R tau was incubated with plasma, it was cleaved into multiple fragments over 48 hours mirroring direct thrombin tau cleavage (**Fig. 7B**). This fragmentation was significantly reduced by PPACK, although some tau digestion products remained, indicating that thrombin alone is not solely responsible for tau degradation. In support of this hypothesis, other M1-associated proteases, such as plasminogen and the metalloproteinase ADAM10, also demonstrated direct tau cleavage (**Supplemental Fig. 6**). Tau degradation was completely blocked only after the addition of a broad-spectrum protease inhibitor cocktail HALT (i.e., AEBSF, Aprotinin, Bestatin, E-64, Leupeptin, and Pepstatin A), confirming that the fragmentation process was driven by protease activity. Consistent with this observation, substitution of plasma with an equivalent concentration of human serum albumin (HSA) did not result in tau cleavage (**Fig. 7A**). Replication of this experiment using pooled Subtype 3 CSF revealed that the plasma proteases in Subtype 3 CSF were also sufficient to induce tau cleavage, producing similar fragmentation patterns to those observed in neat plasma alone (**Fig. 7C**). These findings suggest that elevated plasma protease levels found in M1 in the CSF of Subtype 3 participants, likely due to BBB dysfunction, contribute to tau degradation, providing a potential mechanistic explanation for the lower tau levels observed in this subtype.

**Figure 7:**
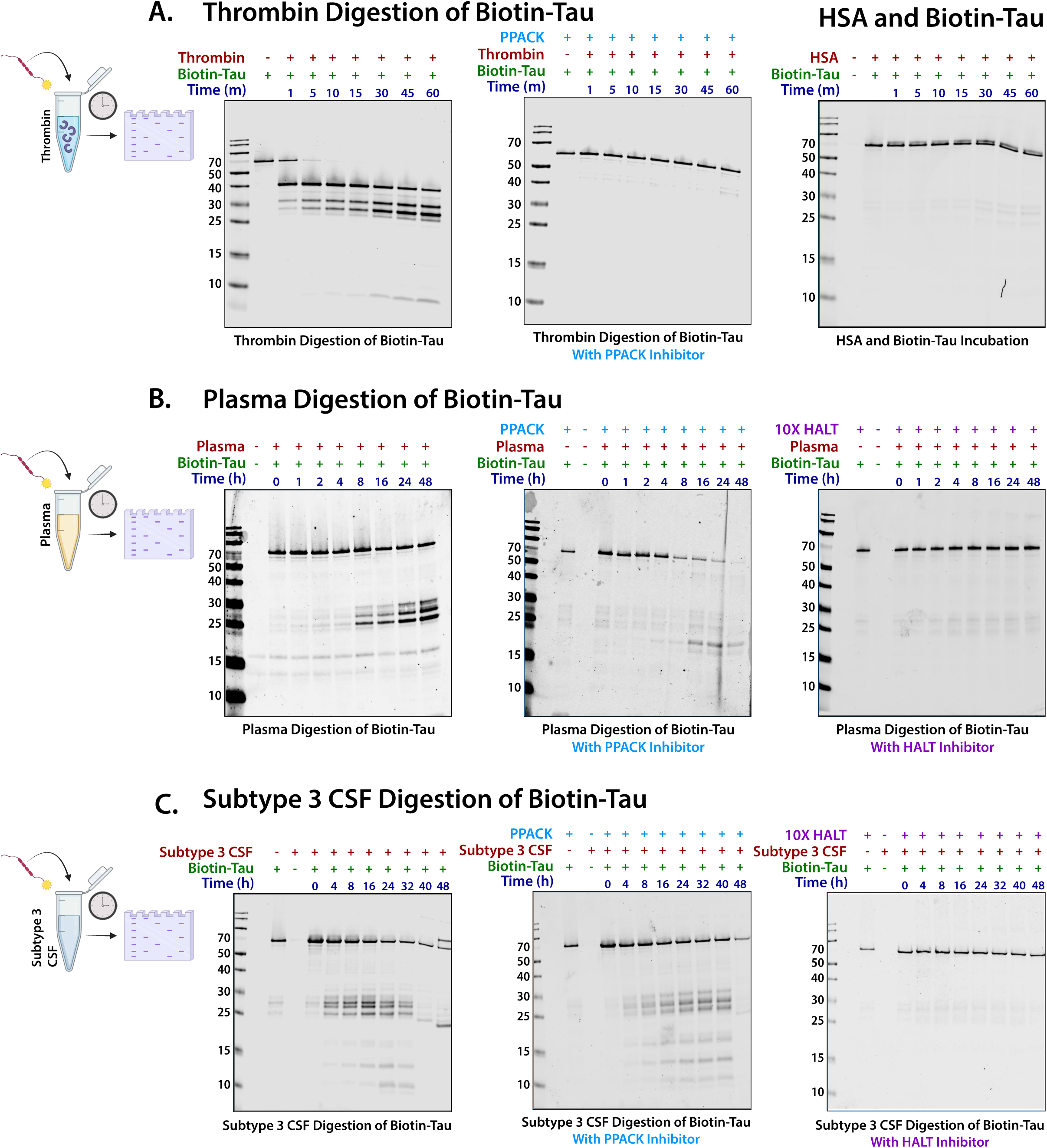
Pooled Plasma and Subtype 3 CSF are capable of cleaving recombinant biotinylated tau. **(A)** Western blot demonstrating thrombin’s cleavage of biotinylated recombinant tau (∼64 kDa) incubated over 60 minutes (**left**) which was prevented by the thrombin specific inhibitor PPACK (**center**). The addition of Human Serum Albumin did not cause Tau cleavage (**right**). **(B)** Pooled human plasma was capable of cleaving recombinant tau over 48 hours (**left**), and a decrease in the production of primary cleavage products (25-27 kDa) at 48 hours was observed when samples were treated with PPACK (**center**), a selective and irreversible thrombin inhibitor. Cleavage was almost entirely inhibited over the course of 48h with the addition of 10x HALT protease inhibitor (**right**). **(C)** Pooled Subtype 3 CSF was also capable of cleaving tau, but cleavage was reduced and delayed with the addition of thrombin inhibitor PPACK (**center**) and militated by 10X HALT protease inhibitor cocktail (**right**).

### Plasma Titration Drives Proteomic Shifts from AD-like Subtype 6 (Neuronal hyperplasticity) to Subtype 3 (BBB Dysfunction)

The strong negative correlations between M1 (BBB/Plasma) and the Cluster III neuronal modules (M2, M8, and M6) suggest that plasma protease activity affects the broader CSF proteome beyond tau degradation. To investigate this, we modeled the biochemical impact of BBB dysfunction by introducing plasma into pooled CSF from Subtype 6, the most prototypical AD-like subtype based on established Aβ and tau biomarker levels. This subtype closely aligns with the neuronal hyperplasticity AD subtype identified in the ACA cohort and is distinguished by its high tau levels and low M1 levels (**Figs. 3** and **5**). This experiment aimed to recapitulate *ex vivo* the increased levels of plasma proteins and proteases observed due to BBB-dysfunction.

Plasma was added to pooled AD-like Subtype 6 CSF at increasing concentrations (0.001%, 0.01%, 0.1%, and 1% by volume) and incubated for 24 hours before analysis by mass spectrometry proteomics (**Fig. 8A**) (**Supplemental Table 17**). As expected, we observed a dose-dependent increase in proteins from M1 (BBB/Plasma) and other blood-associated modules, such as M10 (Peroxidase/Hb), with increasing plasma concentrations in AD CSF. At 1% plasma addition, proteins from M1 increased by 281%, M10 by 126%, and M7 (Extracellular Matrix) by 287% over Subtype 6 baseline (**Fig. 8B-C**). Conversely, we observed a significant reduction in neuronal module proteins, with Cluster III neuronal modules decreasing by approximately 25% from baseline (M8: -28%, M5: -26%, M2: -29%, M6: -24%). Individual proteins exhibited even greater reductions, with key AD-related markers such as SMOC1, MAPT, and GAP43 decreasing by 40–50%, and neuroprotective proteins such as VGF and NRN1 decreasing by nearly 30% (**Fig. 8D-E**). Additionally, Cluster II microglial modules (M3: Collagen, M9: Lysosome/Myelination) showed similar declines (∼25%). Interestingly, M4 (Ubiquitination), which remains stable in AD individuals mapping to Subtype 3 (**Supplemental Fig. 7**), appeared resistant to protease-mediated depletion. The subtyping hub proteins within M4 showed only a 2% decrease from baseline, suggesting protection from degradation. One possible explanation is that M4 is enriched with exosomal proteins (e.g., PPIA, ENO1, PGK1, CFL1, 14-3-3 family), which may be shielded from proteolytic cleavage due to exosome lipid bilayer compartmentalization^48^. As a negative control, we performed the dilution assay and MS-based proteomics using increasing concentrations of human serum albumin (HSA) matching the plasma concentration dilution series, which did not result in a significant decrease in MAPT or members of Cluster III neuronal modules (**Supplemental Fig. 8A**)(**Supplemental Table 18**). Notably, using global protease inhibitors (i.e., HALT) to block enzyme activity precluded the ability to digest the proteins into peptides for MS analysis.

**Figure 8:**
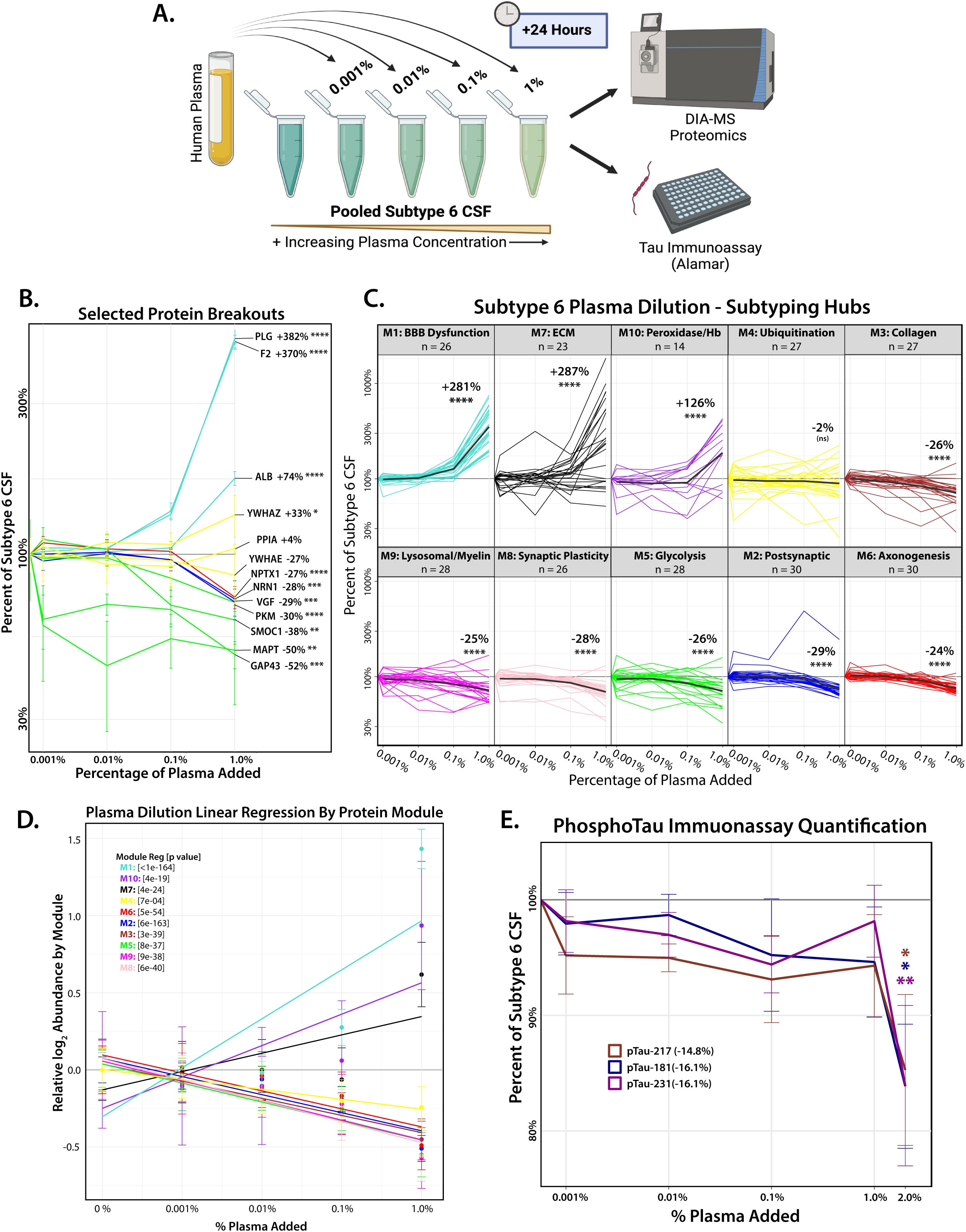
Plasma Dilution into AD-like Subtype 6 CSF Alters Protein Module Abundance and Reduces Tau and Synaptic Protein Levels. **(A)** Schematic for Subtype 6 CSF plasma dilution experiment: a small percentage of human plasma (by volume: 0.001%, 0.01%, 0.1%, 1%) was added to pooled Subtype 6 CSF and incubated for 24 hours. CSF samples were then analyzed by DIA-MS and Alamar immunoassay (with additional 2% plasma concentration) to understand the changes in protein module abundance and immunoassay Tau compared to Subtype 6. **(B)** DIA-MS was used to quantify the proteins present in the samples following plasma dilution. Selected proteins (as a percentage of their baseline at Subtype 6) are broken out to demonstrate the variation in depletion between modules implicated in neurodegeneration and AD pathology (green/yellow), or neuroprotection (blue/red), in response to increasing concentrations of plasma proteases (turquoise). Missing values were imputed, and significance was assessed for the baseline to final concentration by Tukey corrected 1-way ANOVA. **(C)** The proteins that overlapped with the top 30 network hub proteins used to subtype the Emory Cohort were broken out by module, and plotted across increasing amounts of plasma, normalized as a percentage of their abundance in pooled Subtype 6. Bold black lines represent the average percent decrease across the subtyping hub proteins for each module. Significance was determined by Tukey adjusted 1-way rm ANOVA comparing initial and final concentration in each module. One outlier in the blue module (FAM174A) was removed for statistical analysis. **(D)** Linear regression of all proteins quantified by DIA-MS (log_2_ normalized abundance) from each network module, demonstrating the proteome wide trends in response to increasing plasma addition. **(E)** Levels of phosphorylated tau (pTau_181_, pTau_217_, and pTau_231_) present in Subtype 6 CSF was analyzed by Alamar immunoassay following a 24-hour incubation, with increasing concentrations of plasma (by volume: 0.001%, 0.01%, 0.1%, 1%, 2%). This resulted in a significant decrease (∼15%) in all species of pTau. Significance was determined for the final concentration by 1-way ANOVA; (*p≤0.05; **p≤0.01; ***p≤0.001, ****p≤0.0001).

Since peptides corresponding to pTau species were not detected by MS following plasma dilution, we utilized the Alamar NuLISA multiplex immunoassay^49^ to measure endogenous pTau_181_, pTau_217_, and pTau_231_ in AD-like Subtype 6 CSF after dilution with increasing plasma concentrations. Notably, a significant decrease in pTau species was observed at a higher 2% plasma dilution into CSF, where pTau_181_, pTau_217_, pTau_231_ levels in Subtype 6 CSF were reduced by ∼15% from baseline (**Fig. 8F**). In contrast, dilution with an equivalent concentration of HSA had no impact on pTau levels (**Supplemental Fig. 8B**). The higher plasma concentration required to observe a reduction in pTau species may be due to platform differences (MS vs. immunoassay) or the possibility that phosphorylation partially inhibits thrombin cleavage of tau, as previously described^47^. Overall, these findings align with proteome-wide shifts that would occur during a transition from Subtype 6 to Subtype 3, supporting a role of BBB dysfunction and plasma protease activity in shaping distinct CSF biomarkers profiles in AD.

## Discussion

This study highlights the significant molecular heterogeneity of AD by integrating a large, demographically diverse CSF proteomic dataset and identifying distinct biological subtypes. We show that established AD biomarkers, particularly tau, vary by both race and sex, with African Americans, especially men, exhibiting lower CSF tau levels despite comparable Aβ levels to Non-Hispanic White participants. Network-based analysis revealed that these demographic differences extend to broader proteomic patterns, with module M1 (BBB/Plasma) proteins elevated in African Americans and men, while neuronal modules (M5: Glycolysis, M2: Postsynaptic) were higher in women and Non-Hispanic Whites. Using an unbiased clustering approach, we identified six CSF proteomic subtypes, including three AD-like subtypes, two control-like subtypes, and an atypical A+/T- subtype, which had elevated plasma proteases, and reduced neuronal proteins. This subtype, enriched with African Americans and men, was confirmed to have higher CSF/serum albumin ratios in an independent cohort, reinforcing the link between tau depletion and BBB dysfunction. We demonstrate that thrombin and other proteases enriched in M1 can directly cleave tau. Additionally, plasma dilution into the prototypical AD-like Subtype 6 CSF phenocopies the reduced synaptic proteins observed in the atypical A+/T- subtype, leading to tau and neuronal protein depletion, with thrombin, in part, identified as an active contributor to tau cleavage. These findings suggest that BBB dysfunction and plasma protease activity may contribute to lower CSF tau levels, raising concerns about the universal reliability of CSF tau as a biomarker in African Americans, men, and individuals with BBB impairment.

Our findings strongly support and replicate previous subtyping analyses by Tijms and colleagues^25, 26^, as we developed an independent, network-driven molecular subtyping framework that identified six reproducible CSF subtypes. The presence of these subtypes was validated in independent cohorts, including the Swiss dataset^43^, which confirmed the low tau signature of our Subtype 3 and its link to BBB dysfunction through elevated CSF/serum albumin ratios. By demonstrating the reproducibility of these subtypes across diverse cohorts and linking them to underlying biological mechanisms, our study also underscores the importance of demographic-stratified analyses in AD research. For example, although the BBB dysfunction CSF subtype (Subtype 3) is not exclusive to African American men and has been observed predominantly white European cohorts, the inclusion of 111 African Americans and a high proportion of men in this study further accentuated its distinct biological signature, highlighting the intersecting influences of race and sex on AD heterogeneity.

Plasma dilution experiments using AD-like Subtype 6 CSF demonstrated that introducing plasma into neuronal hyperplastic AD subtypes^50^ with high tau and synaptic protein biomarkers (NRGN, GAP43, BASP1, etc.) mirrors the proteomic network module shifts towards Subtype 3, leading to reductions in neuronal proteins, including tau, while increasing plasma-associated proteins. These findings provide a potential explanation for the frequently observed A+T- CSF profile in cognitively impaired individuals, which has raised debate about whether such cases should be classified as AD^42^, given that the biological definition of AD requires abnormal levels of both amyloid and tau^3^. However, biomarkers and their cutoffs may not always accurately reflect underlying pathology. Studies analyzing paired antemortem CSF and brain tissue samples have shown that 73% of A+T- individuals, as determined by CSF immunoassay, were pathologically confirmed to have AD, with both amyloid plaques and tau tangles present at autopsy^42^. Similarly, while lower CSF tau levels have been reported in African Americans, postmortem proteomic studies have found no significant race-related differences in amyloid and tau protein levels in AD brain^8^. Subtype 3 has high levels of neurofilament and other M4 (Ubiquitination) members, measured by proteomics. Collectively, this would suggest that CSF Subtype 3 members with AD have a higher burden of neurodegeneration (N) despite low CSF Tau levels. Given the variability in biomarker cutoffs and the potential for misclassification, particularly in African Americans, our findings underscore the need to account for comorbidities such as blood-brain barrier dysfunction, which may influence CSF tau levels and impact both AD diagnostics and the assessment of tau-targeting therapies in clinical trials. Notably, markers in M4 including 14-3-3 proteins (YWHAZ, YWHAG, YWHAE) were not vulnerable to the effects of plasma dilution into AD-like Subtype 6, possibly due to their enrichment in exosomes^48^. Previously we have reported that these proteins were not influenced by race in the CSF proteome^30^. Given the higher degree of blood brain barrier dysfunction in African Americans it is likely that YWHAZ and other proteins in M4 are more reliable biomarkers of neurodegeneration than total tau itself.

Subtype 3 is notably enriched in African Americans and men, two groups with a higher prevalence of comorbidities such as hypertension and cardiovascular disease^51^, both of which are associated with increased BBB permeability and dementia risk^52, 53^. Thus, further studies incorporating additional patient phenotypes from diverse cohorts are needed to determine whether these comorbidities contribute to this subtype and to assess their impact on disease prognosis. Brain imaging via MRI could help assess microbleeds or infarcts, which may have contributed to the distinct molecular profile of this subtype. Notably, Tijms and colleagues reported that participants in their BBB dysfunction subtype exhibited a higher prevalence of microbleeds on MRI compared to both controls and other AD subtypes^26^, supporting the presence of underlying vascular defects in these individuals.

Tau levels in CSF are likely influenced by multiple processes, including neuronal release, degradation, clearance, and BBB permeability^50^. While plasma proteases may contribute to tau degradation in the CSF, tau is continuously produced by neurons and other cell types, and the overall concentration reflects the dynamic balance between these opposing processes^54^. The reasons why plasma tau, particularly phosphorylated tau (pTau_217_, pTau_231_ and pTau_181_), remains a robust biomarker for AD despite potential for proteolytic cleavage are not fully understood. However, our plasma dilution experiments into AD-like Subtype 6 CSF suggest that phosphorylated tau is partially protected from protease cleavage, supporting previous studies showing that GSK3β phosphorylation inhibits thrombin proteolysis, which cleaves at basic arginine residues^47, 55^. Additionally, tau paired helical filaments from AD brains exhibit increased resistance to thrombin cleavage, an effect reversed by dephosphorylation^47^. It is also important to note that plasma tau and CSF tau levels show poor correlation from the same individuals^56–58^, suggesting that distinct mechanisms and potentially enzymes regulate tau dynamics in these discrete compartments. Despite these complexities, changes in both CSF and plasma ptau still correlate with AD progression, highlighting its utility as biomarker even in the presence of degradation processes.

While our study further supports a subtyping approach to better understand AD heterogeneity, key questions remain. The molecular differences between subtypes may reflect distinct disease etiologies, stages of progression, or comorbid conditions, necessitating further investigation with longitudinal data, imaging, and genetic analyses. Additionally, the clinical implications of BBB dysfunction in AD remain unclear and further studies are needed to assess its impact on disease progression and treatment response. Ultimately, molecular subtyping provides a framework for refining biomarker strategies, improving diagnostic accuracy, and guiding personalized therapeutic interventions that account for the complex and diverse nature of AD pathophysiology.

## Materials and Methods

### Materials

Primary antibodies used in this study were Streptavidin-680 Antibody (Invitrogen, S32358), mouse monoclonal anti-prothrombin antibody (Abcam, ab17199). The secondary antibodies used were conjugated with Alexa Fluor 680.

### CSF Sample Collection and TMT-MS Processing

All CSF samples utilized in this study were collected under the purview of the Goizueta Alzheimer’s Disease Research Center (ADRC), at Emory University. CSF samples were processed in two sets, with sample preparation methods for Set 1 and Set 2 as previously described^29, 30^. Sets 1 and 2 were also independently assessed by immunoassay for diagnostic AD biomarkers such as amyloid beta (Aβ), total Tau, and pTau_181_, with Set 1 run assessed by INNO-BIA AlzBio3 Luminex assay and Set 2 run on the Roche Diagnostics Elecsys platform, as previously described^30^. Z-scoring of each set was performed to normalize these diagnostic biomarkers.

### Database Search and Protein Quantification of CSF Samples

Raw files from Set 1 and Set 2 were downloaded from Synapse and analyzed using the Fragpipe Computational Suite (FragPipe version 18.0, MSFragger version 3.5, Philosopher version 4.4.0, Nesvizhskii Lab). The MS/MS spectra from both sets comprised 1824 raw files across 38 TMT plexes. These sample sets were searched together against the human UniprotKB database (downloaded in 2019, containing 20,402 sequences, including peptides for APOE2 and APOE4) utilizing the following settings: strict trypsin with semi-enzymatic cleavages with a maximum of two missed cleavages, a minimum peptide length of 7 and a maximum length of 35 peptides, and fixed modifications for carbamidomethylation of cysteine (+57.021460 Da), and TMT tags on lysine residues and peptide N-termini (+304.207146 Da). Variable modifications allowed for oxidation of methionine (+15.994900 Da), N-terminal acylation (+42.010600 Da), and additional TMT tag binding on serine, threonine, or tyrosine (+304.207146 Da). Peptide spectral matches were validated using Philosopher, and further filtered based on a false discovery rate (FDR) threshold of 1%, identifying 71,719 peptides which mapped to 4,277 proteins.

### Data Adjustment for Batch and Set Variance

To account for technical batch variance between and within Set 1 and Set 2, we employed a tunable median polish approach (TAMPOR) on proteins that were present in ≥ 50% of samples (n = 2067). Each set was initially run through the TAMPOR algorithm individually and centered to the Global Internal Standards (GIS) included with each batch. The two sets then underwent a second round of TAMPOR together, utilizing the Non-Hispanic White control cases for normalization purposes. Pathophysiological traits, such as Aβ, total Tau, and pTau_181_, were quantified on different immunoassay platforms (Set 1 = Luminex, Set 2 = Roche Elecys), and were normalized by Z-score, resulting in a strong correlation between Z-score and TMT-MS quantified Tau (cor = 0.79, p = 2.9e-104) (**Fig. 1B**). Of the original 504 CSF samples run on the TMT-MS platform, 21 were excluded due to Montreal Cognitive Assessment (MoCA) scores or immunoassay Tau/Aβ ratios that were incongruent with the subject’s stated diagnosis (a Tau/Aβ ratio ≥ 0.226 for was required for confirmation of AD diagnosis), resulting in 483 participant CSF samples. Bootstrap regression was performed to remove any remaining variance within the data due to TMT batch or set, which was confirmed by variance partition analysis (**Supplemental Table 5**), with missing protein values imputed. In both Set 1 and Set 2, samples originating from the same participant’s CSF draw, referred to as “same-case sample-pairs,” were analyzed, resulting in 52 technical replicate pairs. In these cases, only the values from Set 1 were utilized for statistical purposes and represented in participant subtyping.

### Weighted Gene Co-Expression Network Analysis (WGCNA)

The WGCNA algorithm was employed as previously described^28, 30, 34, 59, 60^ to generate a protein co-expression network, across the 483 CSF samples comprised of 245 controls and 238 AD cases, utilizing the following parameters: soft threshold power = 6, deepSplit = 2, minimum module size = 10, merge cut height = 0.7. The resulting network, built on 2067 proteins, was comprised of 10 modules ranging in size from 438 proteins (M1) to 47 proteins (M10), with 489 proteins unassigned to a co-expression module (grey). Correlations between module eigenprotein and demographic and clinicopathological traits were calculated after removing the “same-case sample-pairs” from Set 2 (n = 52). The cell type enrichment of each protein module was determined by Fisher’s exact test, referencing module membership against a list of cell type specific genes, as previously published^30^. Module names, indicative of association to underlying biological processes, were informed by gene ontology (GO) enrichment, utilizing the Bader Lab ontology list, downloaded October, 2020. Bootstrap regression was utilized to isolate protein variance due to sex, race, and diagnosis. Differential expression of these groups was visualized by volcano plot, comparing fold change with Tukey corrected 1-way ANOVA p-value (**Supplemental Tables 10-12**).

### MONET M1 Subtyping

Unbiased proteomic subtyping of the control and AD CSF cases was performed utilizing the MONET M1 modularity network algorithm. To avoid feature redundancy from the larger modules, the top 30 proteins by kME from each module, termed module hubs, were used as the features for a total of 300 proteins across the 10 modules. A grid search was performed, varying minimum module size *i* ∈ *(10, 15, 20, 25),* maximum module size, *j* ∈ *(100, 150, 200, 250, 300, 350, 400)*, and desired average degree, *k* ∈ *(25,50)* ^61^ ^25^. Using a soft power of 13, the following parameters were selected based on performance, (i = 15, j = 200, and k = 25) and produced 6 cohesive subtypes. The six MONET M1 subtypes were generated with the same-case sample-pairs included in the expression data. To assess the model’s accuracy, it was confirmed that same-case sample-pairs were assigned to the same subtype at a rate of 88.5% (n = 46 pairs). Same-case sample-pairs from Set 2 and both samples from ambiguously assigned participant cases were removed from the heat map and subsequent statistical analysis.

### Swiss Cohort UMAP Validation

A supervised UMAP embedding was generated for the 425 Emory cases using the module hub proteins that overlapped with the replication cohort (Dayon et al., n = 167) as features, and the six MONET M1 subtypes as target labels. The supervised dimensionality reduction was performed using UMAP (umap-learn v0.5.4) in Python (v3.11.6) with the following settings: n_neighbors = 10, n_components = 2, metric= Euclidean, and min_dist = 0.1. The resultant clusters are like those produced using MONET M1. Samples used in the embedding were plotted on the reduced dimensions and colored by their original subtype assignment. Five of the 425 samples (1.2%) did not cluster with their previously assigned class. After generating the target clusters, 120 cases from a previously published proteomic study by Dayon et al. were projected onto the UMAP dimension 1 and UMAP dimension 2 space (**Fig. 5B**). Projected cases were assigned to a class by determining which cluster the case was closest to in Euclidean space. If the case was >3 distance away from all clusters, it was determined to be unassigned (n = 5).

### Comparison to the Alzheimer Center Amsterdam Cohort

To compare these Emory Cohort subtyping results to a recently published manuscript on AD subtypes by Tijms, et al., the mean expression of each protein was calculated for each of the six subtypes. As the most cognitively normal group in the Emory Study and comprised of predominantly control cases (95%), Subtype 1 was used as the control reference group. The mean protein expression from the reference subtype was subtracted from the remaining five subtypes to mean center the data on cognitively normal individuals. The resultant mean-centered data was z-scored to produce a mean-centered z-score per protein per subtype. These values were then correlated with the results in Tijms, et al. using the R WGCNA bicor function. A heatmap of the correlation results was produced where the size of each block represents the magnitude of the correlation, with larger block indicating a larger correlation value. Red boxes indicate positively correlated subtypes across the two cohorts, while blue represent negatively correlated subtypes.

### Recombinant Biotin-Tau cleavage by active thrombin

Human active thrombin (ab62452) (10 ng/uL) was added into the recombinant biotin 2N4R tau 441 (2 ng/uL) (rPeptide, T-1114-1) in a phosphate buffered saline (PBS) buffer, pH 7.4 (n=3). The reaction mixture was incubated at 37 °C for intervals of 1 min, 5 min, 10 min, 15 min, 30 min, 45 min and 60 min. A second set of samples was also run in the presence of thrombin inhibitor PPACK dichloride (Abcam, ab141451) (1000 ng/uL). The proteolytic activity of thrombin was stopped by adding 4X Laemmli sample buffer (BioRad, 161-0737) and then denatured at 95 °C for 10 min. The samples were run on a 10% Bis-tris Plus gel, then transferred to nitrocellulose stacks (Thermo, IB23002) for western blots. To identify tau fragments upon thrombin cleavage, the nitrocellulose membrane was immunostained by Streptatvidin-680 antibody (1:1000 Dilution) (Invitrogen, S32358), and the blots were visualized using an Odyssey Imager at 680nm.

### Recombinant Biotin-Tau cleavage by active plasminogen and ADAM-10

Human active plasminogen (Abcam, ab92924) (10 ng/uL) was added into the recombinant biotin tau 441 (2 ng/uL) (rPeptide, T-1114-1) in a PBS buffer, pH 7.4. The reaction was incubated at 37 °C for intervals of 1 min, 5 min, 10 min, 15 min, 30 min, 45 min, 60 min, 90 min and 120 min. The proteolytic activity of plasminogen was stopped by adding 4X Laemmli sample buffer (BioRad, 161-0737) and denatured at 95 °C for 10 min. A second set of samples was run in parallel in the presence of the plasminogen inhibitor tranexamic acid (100 ng/uL) (Thermo, 228040500). The samples were run on a 10% Bis-tris Plus gel then transferred onto nitrocellulose stacks (Thermo, IB23002) for western blot using the iBlot 2 system (Invitrogen, IB21001). To identify tau fragments upon plasminogen cleavage, the nitrocellulose membrane was immunostained by Streptatvidin-680 antibody (1:1000 Dilution) (Invitrogen, S32358), and the blot was visualized using a LI-COR Odyssey Imager at 700 nm. This process was repeated under the same conditions to test the cleavage ability of the metalloprotease ADAM-10, (20 ng/uL) (Randd, 936-AD-020) to digest tau.

### Thrombin Enzyme Activity Assay

Thrombin cleavage was assessed with a substrate specific thrombin assay kit (ab234620). To test the activity of each subtype, representative pooled samples were created from the CSF of the top 40 “hub” members of each subtype by kME. The aliquots of pooled CSF samples were stored at -80 °C for further use. Pooled CSF from each subtype was run with and without the addition of PPACK (100 ng/uL) to test the thrombin cleavage activity of each subtype. Briefly, 10 uL of each subtype pool was added into the 96 well transparent plate (n = 3), followed by 90 uL of assay reaction mixture (1X Diluent and Thrombin Substrate) Reaction samples were incubated at 37 °C, and absorbance readings at 405 nm were collected every 30 min for 24 hours.

### Enrichment of Biotin tau from Pooled Plasma by Streptavidin pull-down

To understand the proteolytic cleavage of tau due to the introduction of plasma proteases, we incubated Biotin Tau (2 ng/uL) (R peptide, T-1114-1) with pooled plasma samples obtained from the Emory Biomarker core at different time points starting from 0 hrs, 4 hrs, 8 hrs, 16 hrs, 24 hrs, 32 hrs, 40 hrs, and 48 hrs at 37 °C. After completion of the incubation, streptavidin pull-down was performed using streptavidin beads (Thermo, 88817) as described previously^62^. Briefly, for each sample, 25 uL of streptavidin magnetic beads were added into a Lo-bind Eppendorf tube and washed twice by adding 1 mL of RIPA buffer (50 mM Tris, 150 mM NaCl, 0.1% SDS, 0.5% sodium deoxycholate, 1% Triton X-100) on rotation for 2 min at RT. The beads were incubated with each sample at 4 °C for 1 hr on rotation. The streptavidin magnetic beads were centrifuged briefly, and flowthrough was collected into new 1.5 mL Lo binding Eppendorf tubes after placing on a magnetic stand rack after 1 hour incubation period. The beads were washed with the following series of buffers on rotation at RT: twice with RIPA lysis buffer (1 mL) for 8 min, once with 1 M KCl (1 mL) for 8 min, once with 0.1 M sodium carbonate (Na_2_CO_3_) (1 mL) for ∼10 s, once with 2 M urea (1 mL) in 10 mM Tris-HCl (pH 8.0) for ∼10 s, and twice with RIPA lysis buffer (1 mL) for 8 min, respectively. After undergoing a final wash with RIPA buffer, the beads were transferred to a new tube and washed twice with PBS buffer (1 mL). The beads were then resuspended in PBS buffer (25 uL). To verify the pull-down, the beads were boiled with 30 uL of 4X sample Lammeli buffer (Biorad, 1610737) supplemented with dithiothreitol (DTT) (20 mM) and Biotin (2 mM) at 95 °C for 15 mins to elute Biotin-Tau. The eluted samples were run on a 10% Bis-tris gel, and a Western blot was performed against a steptavidin-680 antibody (1:1000 Dilution) (Invitrogen, S32358).

### Proteolytic cleavage of Biotin-Tau in pooled Subtype 3 CSF

To investigate the plasma derived enzymatic cleavage activity in Subtype 3 CSF, we performed proteolytic cleavage of recombinant biotin-tau using an aliquot of pooled Subtype 3 CSF. First, the biotin tau 441 (2 ng/uL) (T-1114-1) was added into Subtype 3 CSF. The above reaction mixture was incubated for 0 hrs, 4 hrs, 8 hrs, 16 hrs, 24 hrs, 32 hrs, 40 hrs, and 48 hrs at 37 °C to visualize time dependent proteolytic activity. This experiment was also run in the presence of 100 ng/uL of the thrombin inhibitor, PPACK dichloride (Abcam, ab141451) and 10X HALT protease inhibitor (Thermo, 78429) with same conditions as above to observe the partial and full inhibition of proteolytic activity. After completion of incubation, the proteolytic activity was stopped by adding 4X Laemmli sample buffer (Bio-Rad, 161-0737) and denatured at 95 °C for 10 min. The samples ran on 10% Bis-tris Plus gel, then transferred to nitrocellulose stacks (IB23002) for western blot. To identify the plasma derived proteolytic activity for tau cleavage, the nitrocellulose membrane was immunostained by Streptatvidin-680 antibody (1:1000 Dilution) (Invitrogen S32358) and visualized using an odyssey imager at 680 nm.

### Plasma and human serum albumin dilution of Subtype 6 CSF

To understand BBB dysfunction/damage, we performed an *ex-vivo* dilution of plasma into pooled Subtype 6 CSF, the most “AD-like” subtype, with the highest levels of immunoassay tau. We added plasma at concentrations of 1%, 0.1%, 0.01%, and 0.001% respectively to pooled Subtype 6 CSF and allowed the samples to incubate at 37 °C for 24 hrs. As a control, we replicated this experiment adding human serum albumin (Sigma, A3782-100MG) in lieu of plasma at dilutions, matching the protein content of the plasma dilution concentrations as assessed by BCA assay (Thermo, 23222, 23224). After 24 hours of incubation, the samples were processed for DIA-MS as described below.

### Mass spectrometry of CSF plasma dilution experiments

Following proteolytic digestion as described in previously^30^, samples following were resuspended in an equal volume of loading buffer (0.1% FA, 0.03% TFA, 1% ACN) and analyzed by liquid chromatography coupled to tandem mass spectrometry. Peptide eluents were separated on custom made fused silica column (15 cm × 150 μM internal diameter (ID) packed with Dr. Maisch 1.5um C18 resin) by a Vanquish Neo (ThermoFisher Scientific). Buffer A was water with 0.1% (vol/vol) formic acid, and buffer B was 80% (vol/vol) acetonitrile in water with 0.1% (vol/vol) formic acid. Elution was performed over a 17.5 min gradient. The gradient was from 1% to 99% solvent B. Peptides were monitored on a Orbitrap Astral mass spectrometer (ThermoFisher Scientific) fitted with a high-field asymmetric waveform ion mobility spectrometry (FAIMS Pro) ion mobility source (ThermoFisher Scientific). One compensation voltage (CV) of -35 was chosen for the FAIMS. Each cycle consisted of one full scan (MS1) was performed with an m/z range of 380-980 at 240,000 resolution at standard settings and as many DIA scans in 0.6 second. The higher energy collision-induced dissociation (HCD) tandem scans were collected at 25% collision energy with isolation windows of 2.0 m/z, an AGC of 500% and a maximum injection time set to 2.5ms. Library-free database searches and protein quantification were performed on the DIA-MS raw files using Spectronaut (version 18.1) with default fully tryptic parameter settings. The search database was identical to that used for the TMT-MS analysis. The raw files and database used in this search are available on Synapse. Searched data was column normalized and proteins with ≤ 50% missingness across samples were removed, and missing values were imputed. Proteins overlapping with those identified in the co-expression network were grouped into modules and normalized to their abundance in baseline pooled Subtype 6 CSF (0% plasma or HSA added).

## Availability of data and materials

Raw mass spectrometry data and pre- and post-processed protein expression data and case traits related to this manuscript are available, and can be found at https://www.synapse.org/Synapse:syn65461849. The results published here are in whole or in part based on data obtained from the AMP-AD Knowledge Portal (https://adknowledgeportal.synapse.org). The AMP-AD Knowledge Portal is a platform for accessing data, analyses and tools generated by the AMP-AD Target Discovery Program and other programs supported by the National Institute on Aging to enable open-science practices and accelerate translational learning. The data, analyses and tools are shared early in the research cycle without a publication embargo on secondary use. Data are available for general research use according to the following requirements for data access and data attribution (https://adknowledgeportal.synapse.org/#/DataAccess/Instructions).

## Acknowledgements

We thank the members of the Seyfried Lab and the Center for Neurodegenerative Disease for their valuable comments and discussions throughout the completion of this manuscript.

## Author Contributions

Conceptualization, MCB, NTS; Methodology, MCB, EKC, LP, DMD, EBD, JG, NTS; Investigation, MCB, JG, LP, WF, DMD; Formal Analysis, MCB, EKC; Writing – Original Draft, MCB and NTS; Writing – Review & Editing; Funding Acquisition, JJL, AIL, and NTS; Resources, JJL, AIL, and NTS, and.; Supervision, JJL, AIL, and NTS. All authors read and approved the final manuscript.

---

**Supplemental Figure 1:**
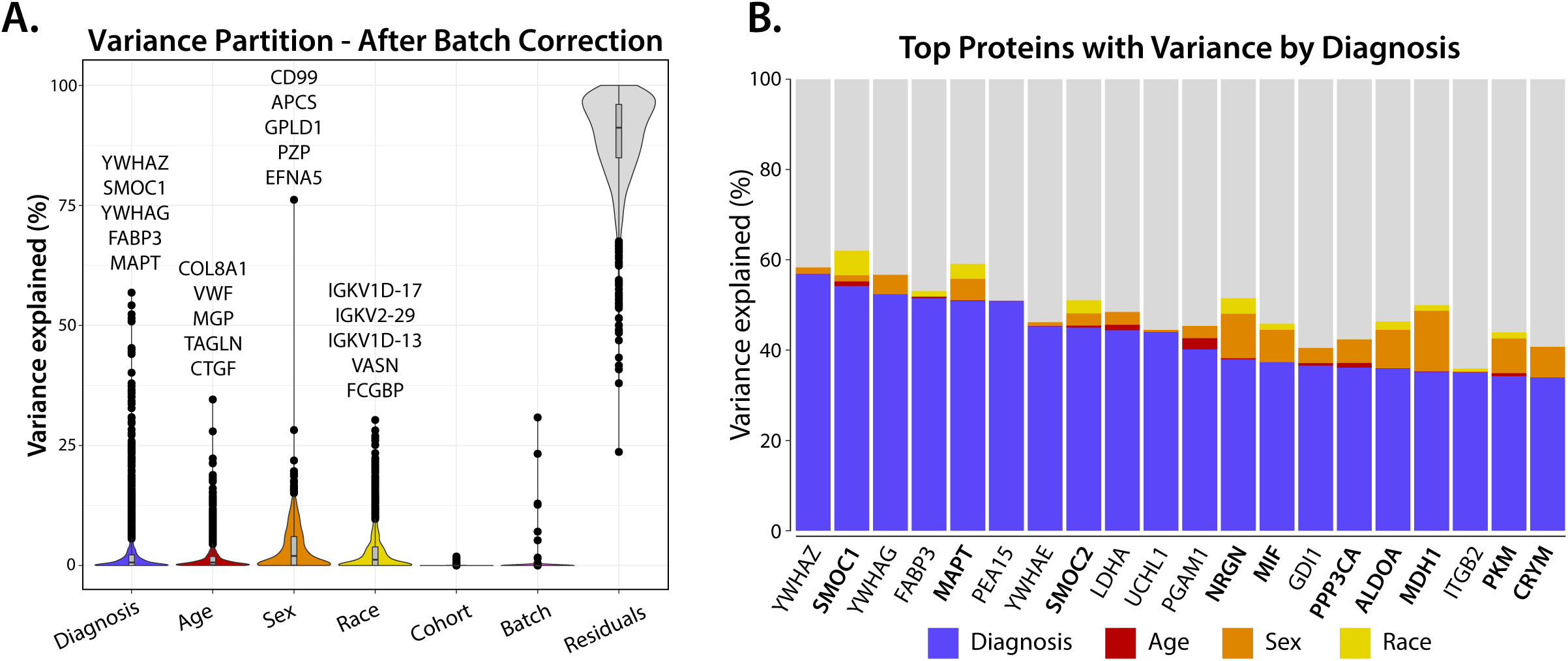
Variance partition analysis reveals that proteins associated with AD diagnosis differ based on race and sex. **(A)** Variance partition analysis was performed to determine to what magnitude the factors of diagnosis, age, race, and sex contributed to variations in cohort protein abundance, and to confirm minimal contributions to data variance due to cohort or TMT batch. The top 5 proteins with the greatest variance in abundance attributable to each factor are noted. **(B)** The top 20 proteins with the greatest variance in abundance across cases due to AD diagnosis also have notable contributions to their variance attributable to the sex, race, and age of the participant. Bolded proteins labels indicate proteins where the combined variance due to race and sex is greater than 10% of that due to diagnosis.

**Supplemental Figure 2:**
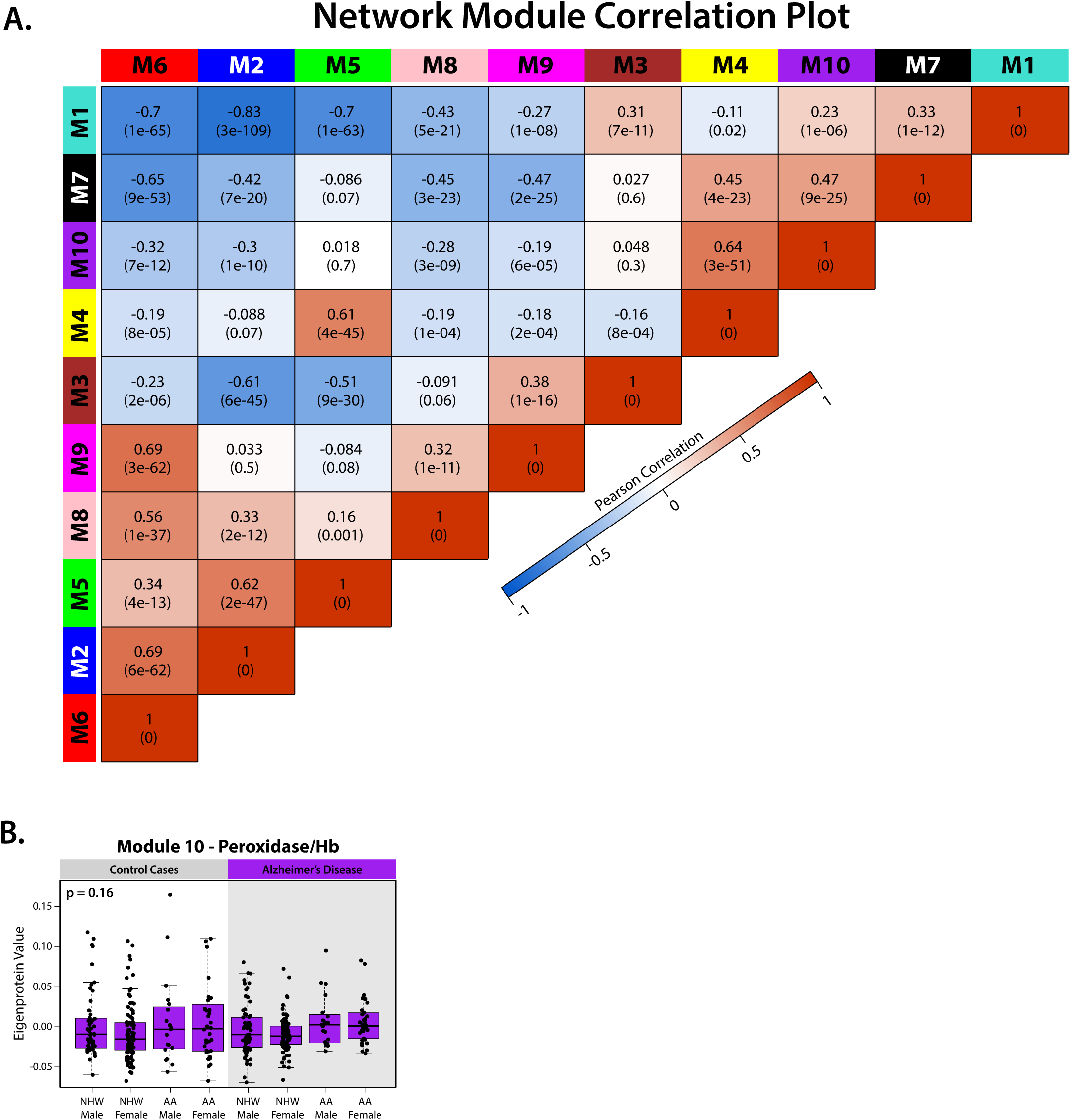
Module correlations within the co-expression network. **(A)** Pearson correlations and p values corresponding to protein modules within the network demonstrating positive (red) or negative (blue) associations. **(B)** Eigenprotein values from Module 10 broken out by participant demographic. Significance assessed by 1-way ANOVA, points outside of 3 standard deviations for each subtype were not plotted.

**Supplemental Figure 3:**
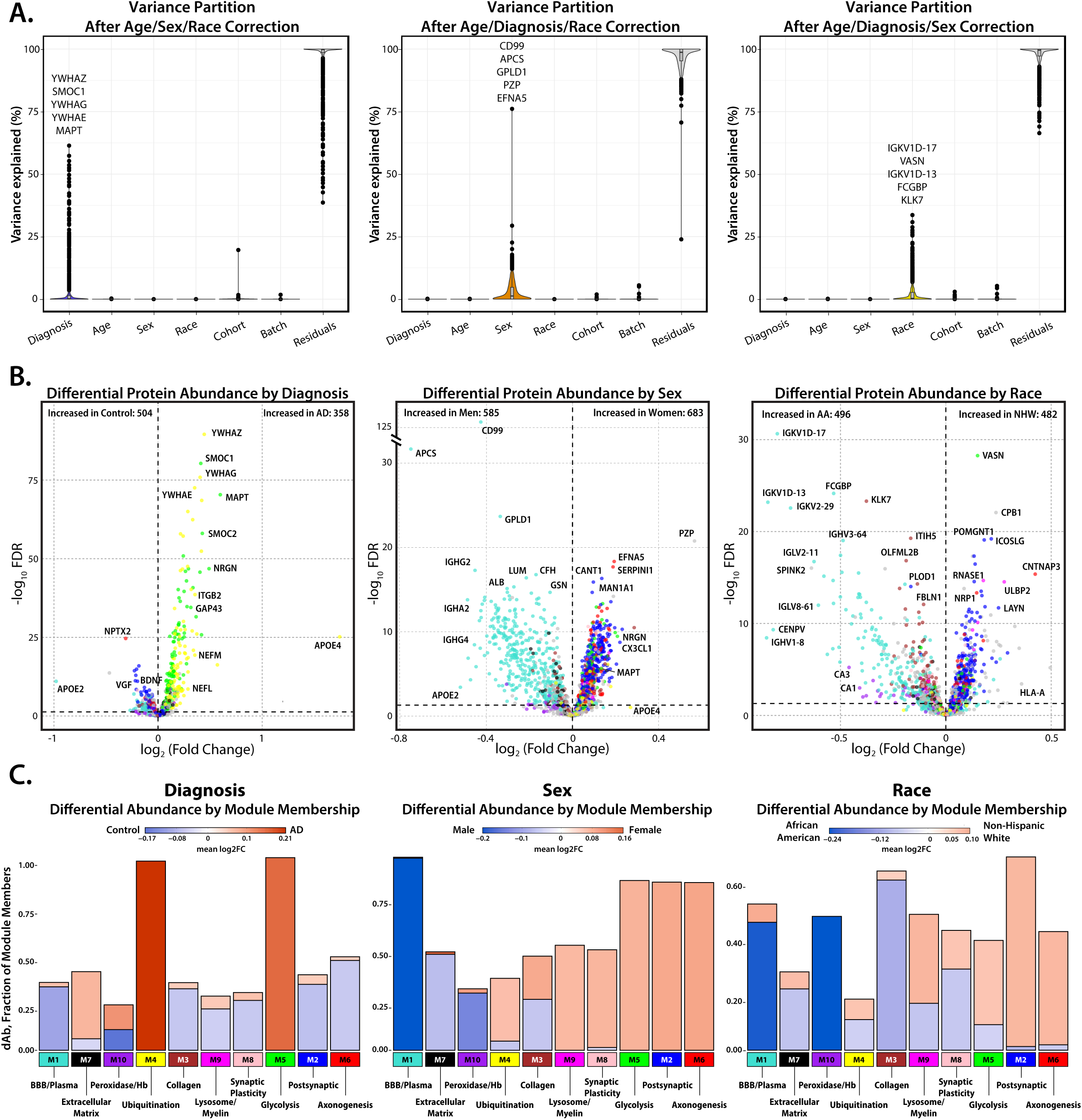
Influence of Diagnosis, Sex, and Race on Protein Variance and Module Abundance in the Emory Cohort. **(A)** Variance partition analysis of the Emory cohort regressed for all variables except diagnosis (**left**), sex (**center**) and race (**right**) demonstrate the effective minimization of other sources of variance. Plots are annotated with the top 5 proteins with the greatest variance in abundance due to each factor. **(B)** Volcano plots displaying the log2-fold change of protein abundance plotted against the FDR corrected 1-Way ANOVA p value, demonstrate the differential protein abundance of the preserved factors of diagnosis (**left**), sex (**center**) and race (**right**) in the absence of other variables. Proteins are annotated by module color, illustrating the relationships between modules and demographic factors. (**C**) A fractional breakdown of the proteins in each module that have significant differential abundance based on the selected factors of diagnosis (**left**, blue: control, red: AD), sex (**center**, blue: male, red: female) and race (**right**, blue: AA, red: NHW), highlighting the fundamental influence of diagnosis and demographics on module abundance, even in the absence of the other sources of variance.

**Supplemental Figure 4:**
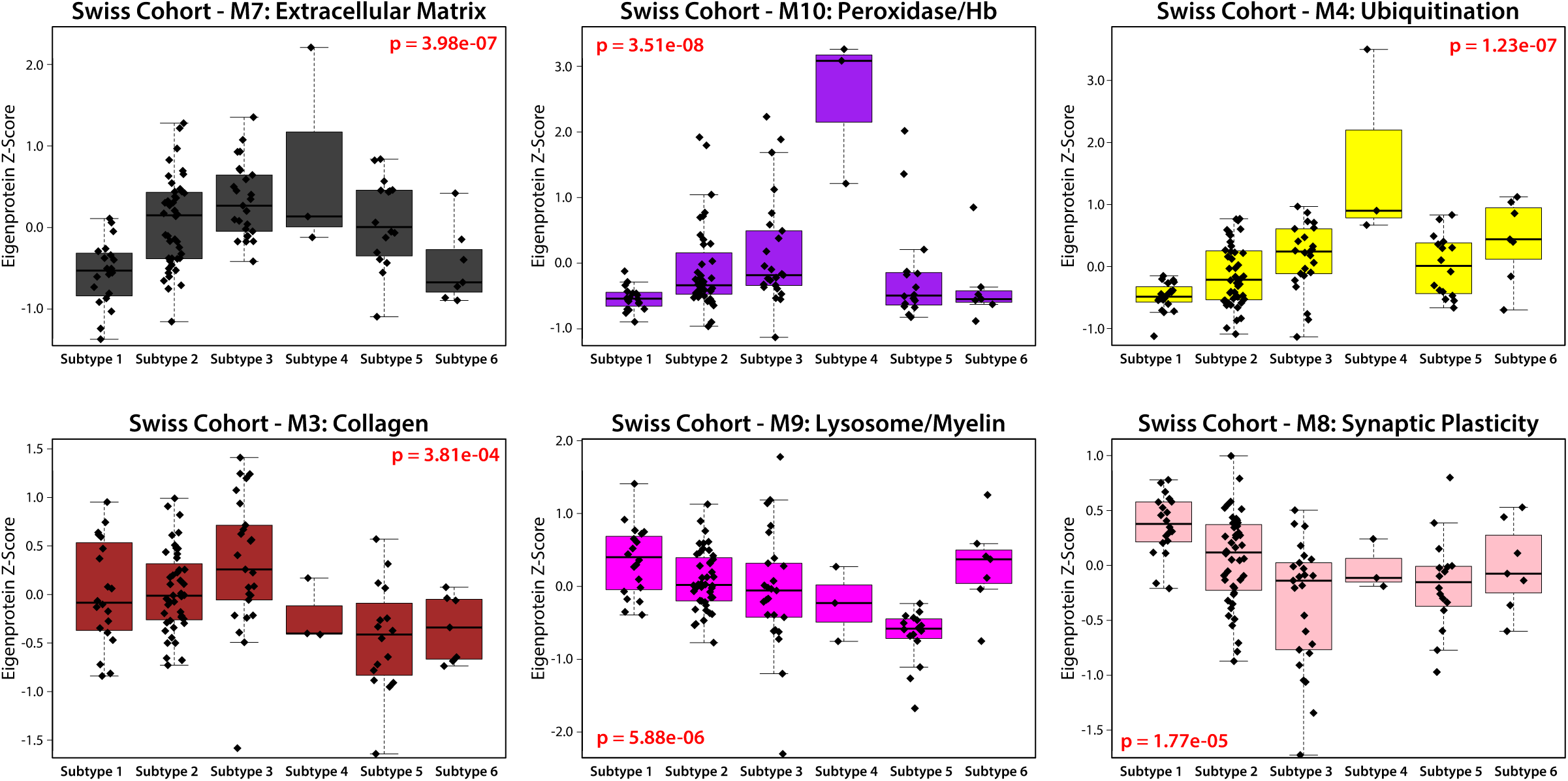
Network Module Eigenprotein Levels Across UMAP Subtypes in the Swiss Replication Cohort. Additional network module eigenprotein boxplots for the Swiss Replication cohort, broken out by assigned UMAP subtype in order of module relatedness. Significance was assessed by 1-way ANOVA, points outside of 3 standard deviations for each subtype were not plotted.

**Supplemental Figure 5:**
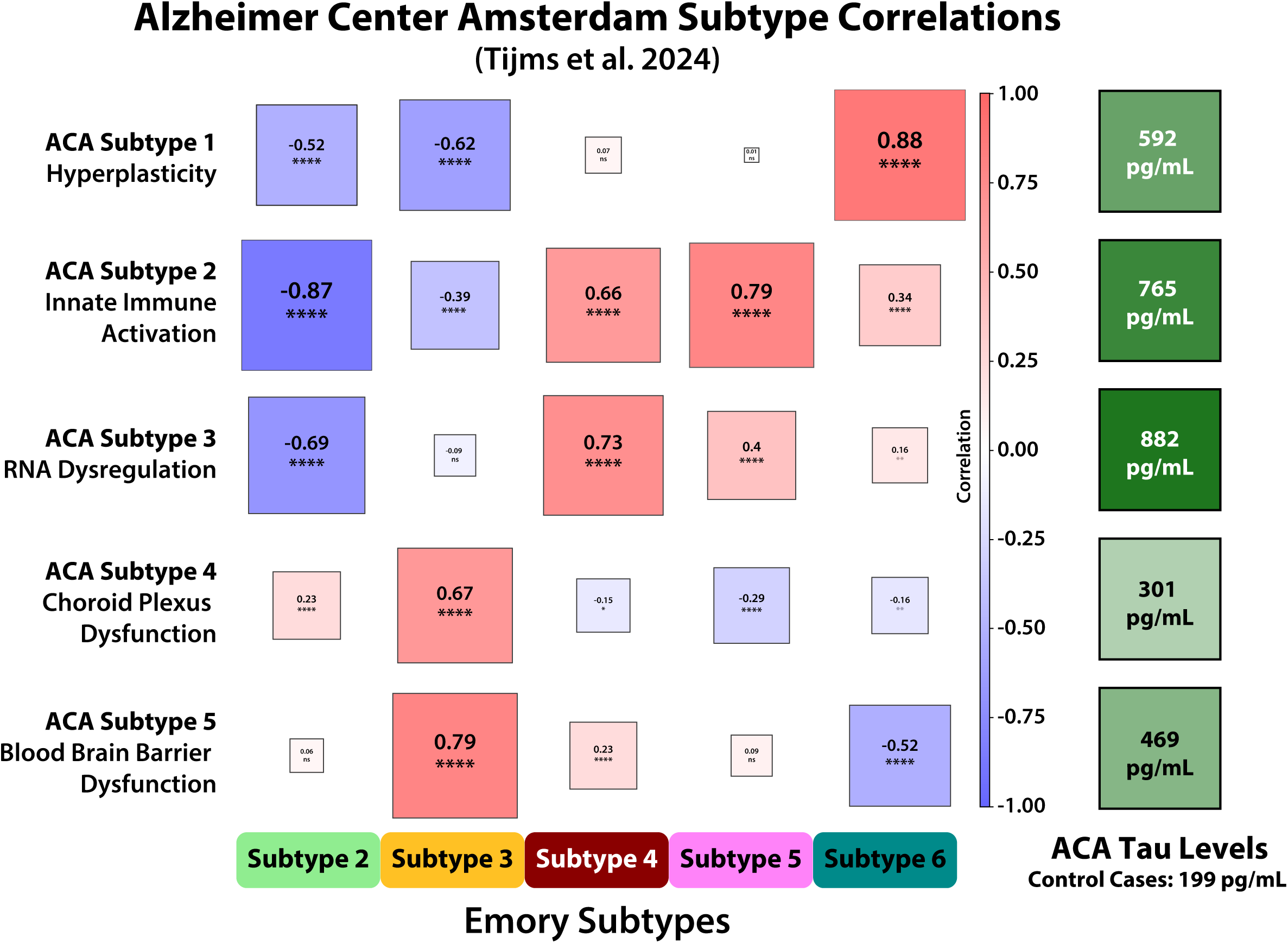
Correlation between the Emory and Alzheimer’s Center Amsterdam Subtypes. **(A)** A heatmap showing the correlation between the z-scored mean protein abundances of participants in the Emory and Alzheimer’s Center Amsterdam (ACA) proteomic subtypes. Subtypes with strong positive (red) or negative (blue) bicor values between the cohorts indicate relatedness (*p≤0.05; **p≤0.01; ***p≤0.001 ****p≤0.0001). **(B)** Corresponding immunoassay CSF Tau levels from each ACA subtype, where all participants have been diagnosed with AD. ACA subtypes that were highly correlated with Emory Subtype 3 (ACA Subtype 4/Choroid Plexus Dysfunction, ACA Subtype 5/Blood Brain Barrier Dysfunction) also had the lowest levels of CSF Tau.

**Supplemental Figure 6:**
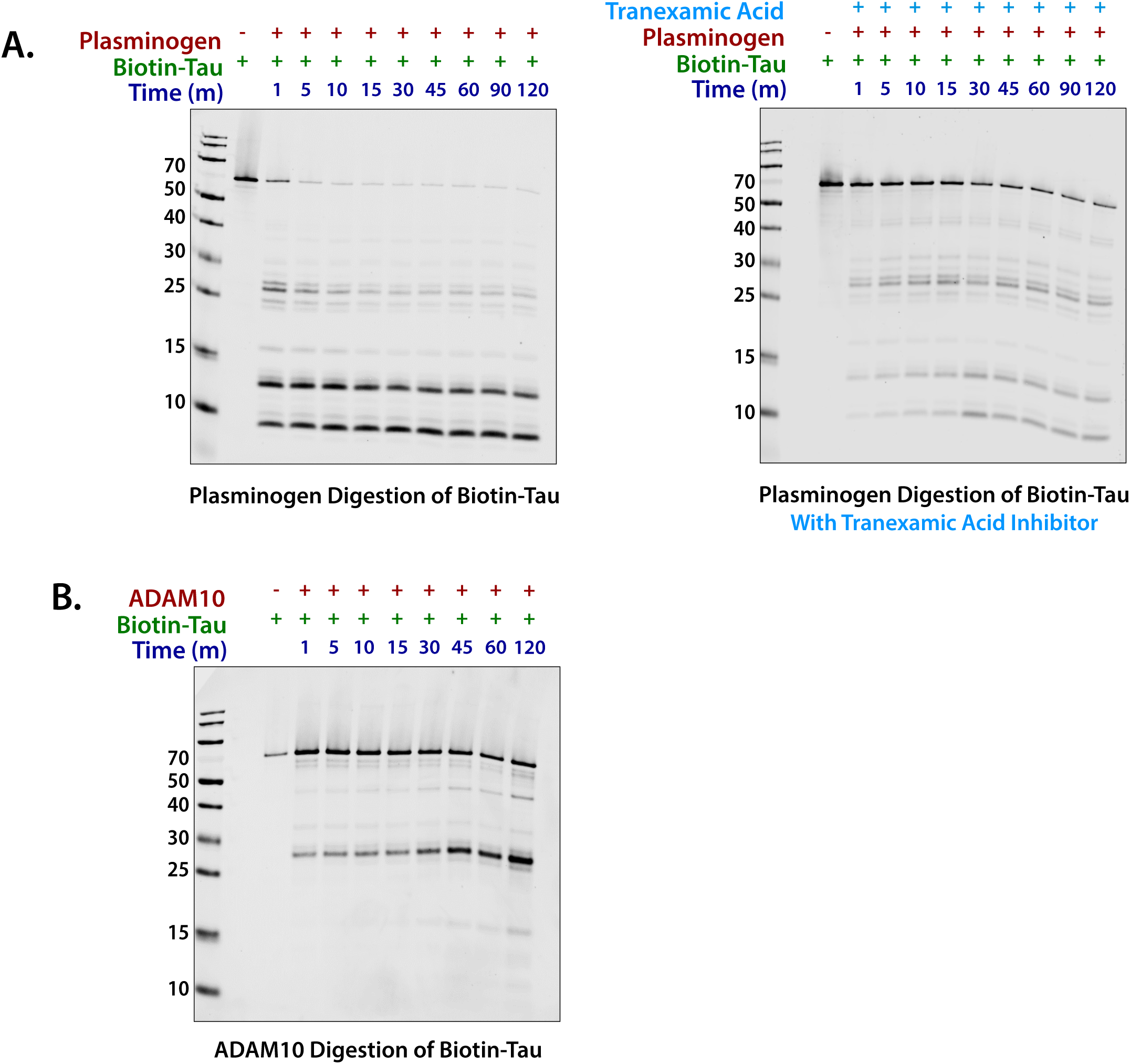
Module 1 members Plasminogen and ADAM10 are capable of cleaving recombinant tau. **(A)** Western blot demonstrating that plasminogen is capable of cleaving biotinylated recombinant tau (∼64 kDa) over 120 minutes, a process which is partially inhibited by tranexamic acid, as is ADAM10 (**B**).

**Supplemental Figure 7:**
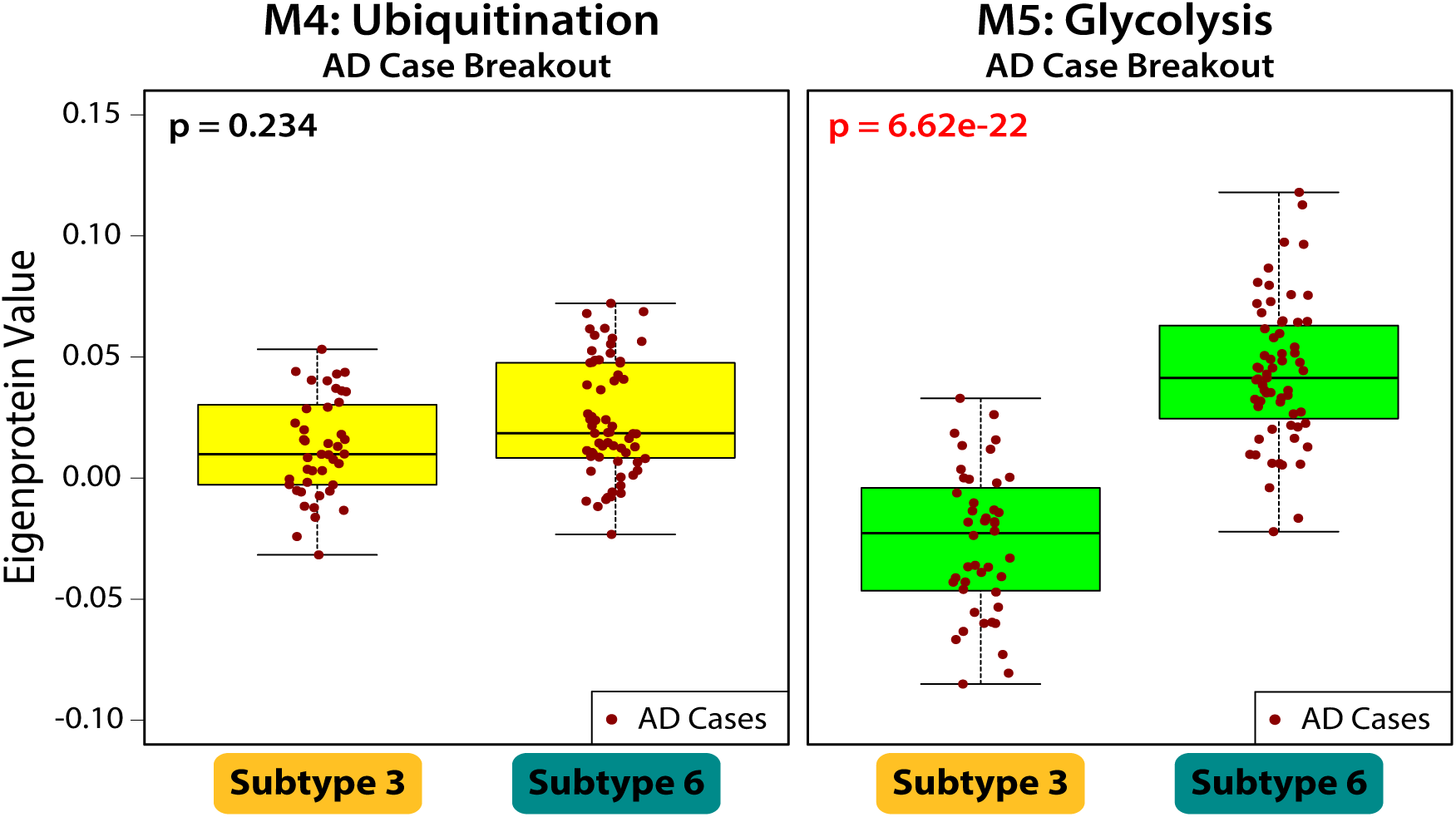
Comparison of AD Pathology-Associated CSF Modules Between Subtype 6 and Subtype 3 AD Samples. A breakout of exclusively the AD cases from Subtype 6 and Subtype 3, comparing levels of the two AD pathology associated modules, M5: Glycolysis, and M4: Ubiquitination. While there is a significant difference in M5 levels in AD cases between the two subtypes, there is no significant difference in M4 levels (as assessed by 1-way ANOVA).

**Supplemental Figure 8:**
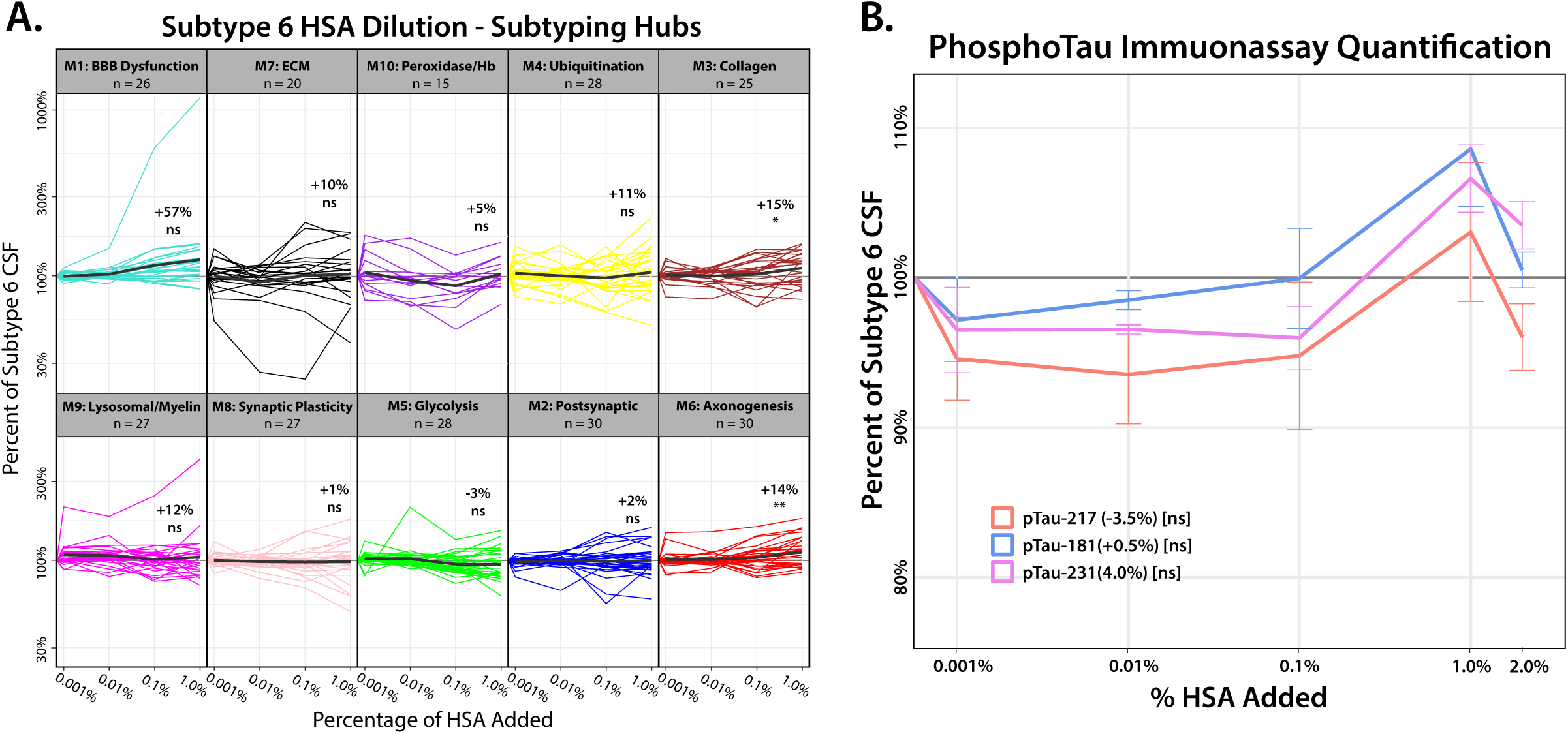
Impact of Human Serum Albumin (HSA) Dilution on Network Module Proteins and Phosphorylated Tau Levels in AD-like Subtype 6 CSF. **(A)** DIA-MS analysis of HSA doping was run in parallel to plasma doping experiments. Proteins that overlapped with the top 30 network hub proteins used to subtype the Emory Cohort were broken out by module, and plotted across increasing amounts of HSA, normalized as a percentage of their abundance in pooled Subtype 6. Bold black lines represent the average percent decrease across the subtyping hub proteins for each module. Significance was determined by Tukey adjusted 1-way rm ANOVA comparing initial and final concentrations in each module. **(B)** Levels of phosphorylated tau (pTau_181_, pTau_217_, and pTau_231_) present in pooled Subtype 6 CSF were analyzed by Alamar immunoassay following a 24-hour incubation, with increasing concentrations of HSA (by volume: 0.001%, 0.01%, 0.1%, 1%, 2%). HSA addition did not significantly decrease endogenous levels of any phosphorylated tau species. Significance was assessed for the final concentration by 1-way ANOVA; (*p≤0.05; **p≤0.01; ***p≤0.001, ****p≤0.0001).

